# The extracellular ATP/P2X7R signaling axis drives early neuroinflammation and neuronal hyperexcitability in an Alzheimer’s disease mouse model

**DOI:** 10.1101/2025.11.14.688405

**Authors:** Nikita Arnst, Martina Bedetta, Nelly Redolfi, Simonetta Falzoni, Neha Kachappilly, Mario Tarantini, Emy Basso, Anna Lisa Giuliani, Dorianna Sandonà, Francesco Di Virgilio, Annamaria Lia, Elisa Greotti, Paola Pizzo

## Abstract

Neuroinflammation and synaptic dysfunction are emerging as early and potentially causative events in Alzheimer’s disease (AD), yet their molecular triggers remain elusive. Here, we identify extracellular ATP (eATP), a major damage-associated molecular pattern, and its purinergic receptor P2X7 (P2X7R) as pivotal drivers of early pathology in AD mice. *In vivo* bioluminescence imaging revealed a significant cortical accumulation of eATP in AD mice as early as 2 months of age—before amyloid plaque deposition and cognitive impairment. This increase is associated with inflammasome activation, pro-inflammatory cytokine production, microglia reactivity, aberrant synaptic pruning and perineuronal net degradation. Strikingly, genetic deletion of P2X7R rescues these alterations. Two-photon calcium imaging further demonstrates that P2X7R knockout counteracts AD-related neuronal hyperactivity. These findings set the eATP–P2X7R signaling axis as an early driver of AD pathology, linking neuroinflammation to synaptic remodeling and circuit dysfunction, and suggest P2X7R inhibition as a compelling strategy to counteract AD progression.

## Main text

Alzheimer’s disease (AD) is a progressive neurodegenerative disorder that begins decades before clinical symptoms. While amyloid-β (Aβ) plaques and hyperphosphorylated tau tangles remain defining histopathological hallmarks, increasing evidence suggests that neuroinflammation — largely driven by reactive microglia — is a central trigger and amplifier of AD^1^.

In the healthy brain, microglia sustain homeostasis by clearing apoptotic debris, pruning synapses, shaping plasticity, regulating neuronal excitability^2^, and remodeling the extracellular matrix (ECM)^3^. In AD, research has mainly focused on the late microglial response to Aβ accumulation, particularly their recruitment around plaques and the adoption of a pro-inflammatory, maladaptive phenotype^4^, which fuels a self-amplifying loop of neuronal damage and network instability^5^. However, synaptic deficits, cortical hyperexcitability and neuroinflammation often precede overt Aβ plaque deposition and memory decline and are now recognized as early functional phenotypes of AD^1^. Although microglia play a central role in synaptic and ECM remodeling and inflammation, their contribution to these possible early alterations in AD remains poorly understood, and the molecular signals linking microglial dysfunction to circuit pathology are still largely undefined^6^. Extracellular ATP (eATP), a prototypical damage-associated molecular pattern (DAMP), is a key signal that initiates and propagates inflammatory responses. Released into the extracellular *milieu* through lytic and non-lytic pathways — including pannexins, connexins, and the purinergic P2X7 receptor (P2X7R) — eATP reaches high concentrations (>100 μM) at sites of trauma, tumors, and inflammation, with similar accumulations also occurring in diseased brains^7^. Once released, eATP is degraded by ectonucleotidases, such as CD39 and CD73, into adenosine (ADO), which blunts inflammation via P1 adenosine receptor activation, before its further breakdown by adenosine deaminase (ADA)^8^. Thus, the dynamic eATP/ADO ratio represents a key regulator of tissue inflammatory balance. Moreover, recent studies identified eATP as a danger signal released by stressed or hyperactive neurons^9^, whereas ADO acts as a neuroimmune modulator that counteracts neuronal hyperexcitability and inflammation^10^.

Among purinergic receptors, eATP primarily signals through P2X7, P2X4, P2Y2, P2Y6, and P2Y12. P2X7R is unique in its low affinity and high activation threshold, requiring high eATP levels typically found at inflammatory sites. Once activated, P2X7R mediates both sodium (Na⁺) and calcium (Ca²⁺) influx, and potassium (K⁺) efflux, and acts as a key activator of the NLRP3 inflammasome. This ultimately leads to IL-1β and IL-18 processing and release, and, in severe cases, to pyroptosis^11^. Uniquely among purinergic receptors, P2X7R, following tonic and prolonged activation, can form a membrane macropore, allowing passage of molecules up to 900 Da, including ATP itself and promoting secretion of cytokines and chemokines (*e.g.*, TNF-α, IL-6, CCL3), as well as the production of reactive oxygen species (ROS)^11^.

Remarkably, P2X7R has been found upregulated in Aβ plaque-associated microglia from both AD patients and mouse models^12,13^. Recent studies suggest that its inhibition alleviates neuroinflammation and cognitive symptoms in different AD mouse models and its expression is essential for Aβ-dependent microglia activation, NLRP3 inflammasome stimulation, IL-1β release, and Aβ-induced microglia dysfunction^14^. However, whether eATP–P2X7R signaling contributes to the initial stages of AD pathogenesis remains unresolved.

Here, we used the B6.152H AD mouse model (hereafter referred to as AD mice), which recapitulates the progressive nature of AD pathology^15^. Combining *in vivo* imaging of cortical eATP, high-resolution microglia morpho-functional analyses, inflammasome and cytokine assays and two-photon (2P) Ca²⁺ imaging in somatosensory cortex (SSCx) brain slices, we identify the eATP–P2X7R pathway as a primary driver of AD onset, preceding Aβ deposition and linking microglia reactivity to circuit dysfunction. Notably, genetic deletion of P2X7R rescues these phenotypes, highlighting P2X7R as a promising therapeutic target within an early disease time-window, before Aβ plaque deposition and irreversible degeneration.

## Results

### eATP-P2X7R signaling triggers early neuroinflammation in AD mice

eATP acts as a danger signal and pro-inflammatory mediator in multiple pathological settings, including AD^7^. However, its direct *in vivo* measurement in the brain has been technically challenging. Using the plasma membrane-targeted luciferase probe pmeLUC, a bioluminescent reporter that allows semi-quantitative detection of eATP in live animals^16^(Fig. 1a), we addressed this issue for the first time in an AD mouse model. Following retro-orbital AAV-pmeLUC delivery, we monitored cortical eATP in AD mice at 2 months (pre-symptomatic stage, before Aβ plaque deposition), 6 months (prodromal stage, initial Aβ plaque appearance), and 9 months (overt Aβ plaque load and cognitive impairment)^15^ (Fig. 1b). As expected, cortical eATP levels were elevated in AD mice compared with wild-type (WT) controls at 6 and 9 months (Fig. 1c,d), coinciding with Aβ plaque deposition and activation of the inflammatory response. Remarkably, however, brain eATP levels were already increased in 2-month-old AD mice (Fig. 1c,d), well before Aβ plaque deposition, suggesting a potential role for this molecule at the pre-symptomatic stage of the disease. Subsequent analyses focused on 2 and 6 months, as this timeframe captures the transition from a pre-plaque condition to the early appearance of Aβ plaques and initial cognitive deficits^17^.

**Figure 1.**
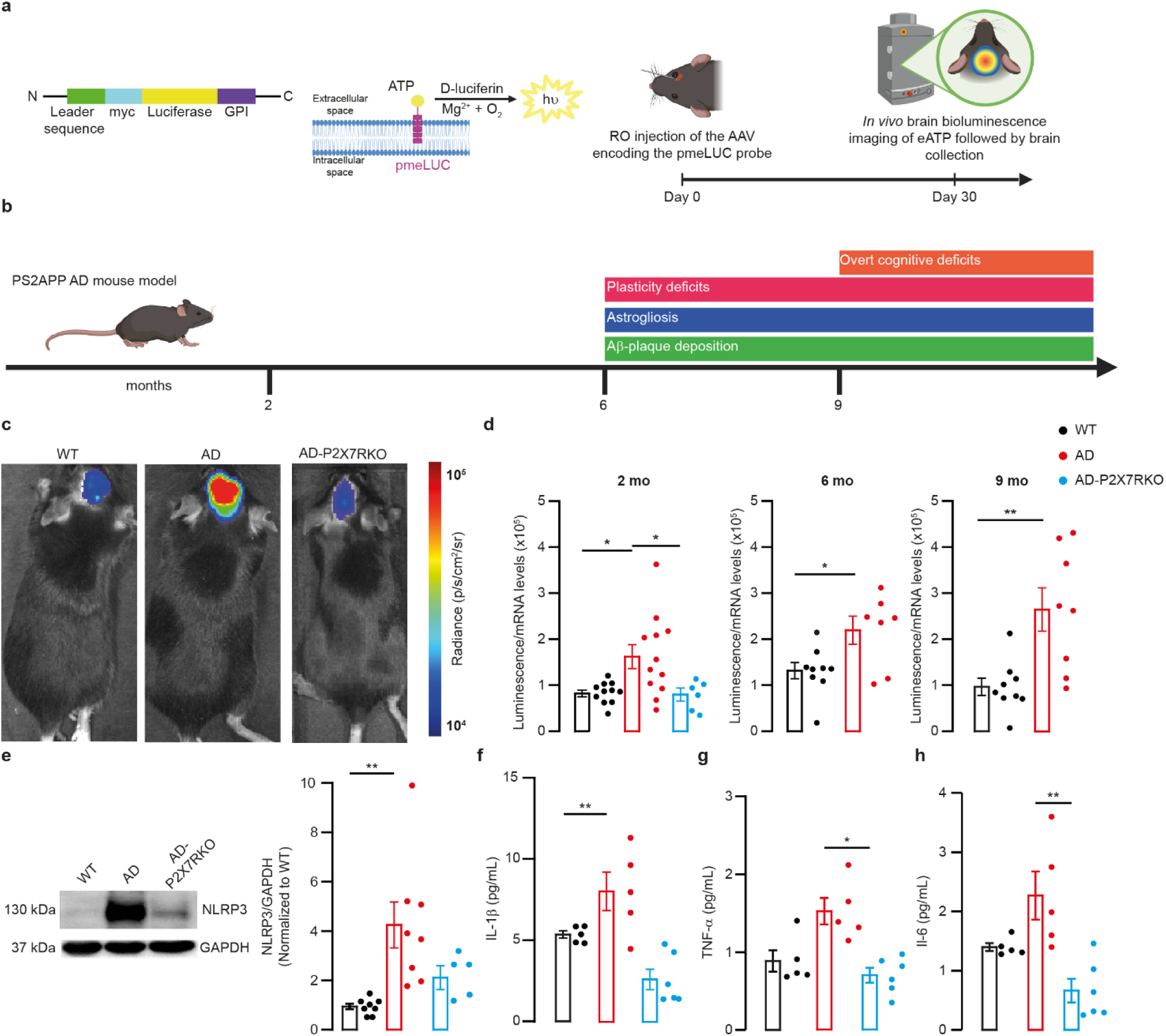
Early cortical eATP accumulation and inflammasome activation in AD mice are P2X7R-dependent. **a,** Left, schematic of the pmeLUC probe functioning. The plasma membrane–anchored luciferase generates light in the presence of ATP and D-luciferin. Right, schematic of the protocol for *in vivo* bioluminescence imaging of cortical eATP using the pmeLUC probe in mice. AAV–pmeLUC was delivered via retro-orbital (RO) injection, and cortical eATP was imaged *in vivo* 30 days later, using an IVIS system. Created with BioRender.com. **b**, Timeline of pathological events in the PS2APP (AD) mouse model showing onset of synaptic plasticity deficits, astrogliosis, Aβ-plaque deposition and cognitive decline. Created with BioRender.com. **c**, Representative IVIS-100 bioluminescence images of 2-month-old WT, AD, and AD-P2X7RKO mice. Color scale indicates radiance (photons/s/cm²/sr). **d**, Quantification of total luminescence values normalized to cortical pmeLUC mRNA levels in WT (black), AD (red), and AD-P2X7RKO (cyan) mice at 2 (2 mo; WT, n = 13; AD, n = 14; AD-P2X7RKO, n = 8), 6 (6 mo; WT, n = 9; AD, n = 7), and 9 (9 mo; WT, n = 9; AD, n = 8) months of age. **e**, Representative immunoblot of NLRP3 protein, with relative quantification, in SSCx lysates from 2-month-old WT (n = 8), AD (n = 8), and AD-P2X7RKO (n = 5) mice. Data are presented as mean ± s.e.m., normalized first to GAPDH and then to WT. **f–h,** Levels of IL-1β (f), TNF-α (g), and IL-6 (h) cytokines in SSCx from 2-month-old WT (n = 5), AD (n = 5), and AD-P2X7RKO (n = 6) mice, measured by ELISA. Data are presented as mean ± s.e.m.; statistical analysis was performed using Kruskal–Wallis test followed by Dunn’s post hoc test. For data collected at 6 and 9 months (d), comparisons between WT and AD mice were performed using unpaired two-tailed t-tests. \**P* < 0.05; \*\**P* < 0.01.

To test whether P2X7R contributes to eATP brain accumulation, we generated AD-P2X7RKO mice by crossing AD with P2X7R knockout (KO)^18^ animals. Strikingly, cortical eATP levels in 2-month-old AD-P2X7RKO mice are indistinguishable from those detected in WT (Fig. 1c,d), indicating that P2X7R is required to sustain early eATP high concentrations in the AD brain.

This early eATP accumulation is functionally linked to NLRP3 inflammasome activation, since cortical NLRP3 protein levels are increased in 2-month-old AD mice compared with both WT and AD-P2X7RKO mice (Fig. 1e). Consistently, cortical IL-1β levels are elevated in 2-month-old AD mice, whereas AD-P2X7RKO animals display levels comparable to WT (Fig. 1f). Similar increases in NLRP3 and IL-1β levels are present in 6-month-old animals (Extended Data Fig. 1b).

Additional cytokine analyses show that at 2 months, cortical TNF-α (Fig. 1g) and IL-6 (Fig. 1h) are also increased in AD, but not in AD-P2X7RKO mice. At 6 months, both TNF-α and IL-6 levels are similar in WT and AD but lower in AD-P2X7RKO mice (Extended Data Fig. 1c,d).

eATP accumulation in the tissue *interstitium* can arise from increased cellular release and/or decreased extracellular degradation. Since ATP release can occur through P2X7R itself, when activated in its pore-forming structure by chronically elevated eATP concentrations^11^, we analyzed P2X7R expression in the brain at different time points. *P2rx7* mRNA expression is unchanged at 2 months but is increased in AD brains at 6 months (Extended Data Fig. 1e), indicating that the earliest eATP accumulation precedes *P2rx7* transcriptional upregulation. At 6 months, instead, *P2rx7* upregulation could contribute to sustaining high eATP levels and likely represents a consequence of Aβ plaque deposition^12^.

As for eATP-degrading pathways, we examined brain expression of ectonucleotidases. *Entpd1* (CD39) mRNA is significantly increased in AD mice only at 6 months (Extended Data Fig. 1f), whereas *Nt5e* (CD73) mRNA remains unchanged compared with WT animals at both time points (Extended Data Fig. 1g), likely suggesting that impaired eATP degradation does not account for its early elevation in the brain.

### P2X7R drives early microglia reactivity in AD mice

Morphological remodeling is a hallmark of microglia activation, reflecting transitions from homeostatic to reactive states that contribute to neurodegeneration^19^. Once microglia become reactive, they undergo morphological changes, adopting a more amoeboid-like shape with altered soma size and process complexity — features typically associated with the loss of physiological functions and the acquisition of pro-inflammatory or neurotoxic phenotypes^20^. While microglia reactivity is well established in Aβ-laden AD brains^21^, it remains unclear whether similar changes occur before Aβ plaque deposition and which mechanisms drive them.

To address this, we analyzed microglia changes across layer IV (L4) of the SSCx in WT, AD, and AD-P2X7RKO mice. At 2 months, microglial density is unchanged across genotypes, indicating that extensive proliferation is negligible at early AD stages (Fig. 2a; Extended Data Fig. 2a). By 6 months, however, microglial density is significantly increased in AD mice compared with WT, an effect not reversed by P2X7R deletion (Extended Data Fig. 2h), suggesting that the receptor does not mediate the proliferative microglial response associated with Aβ plaque deposition and disease progression.

**Figure 2.**
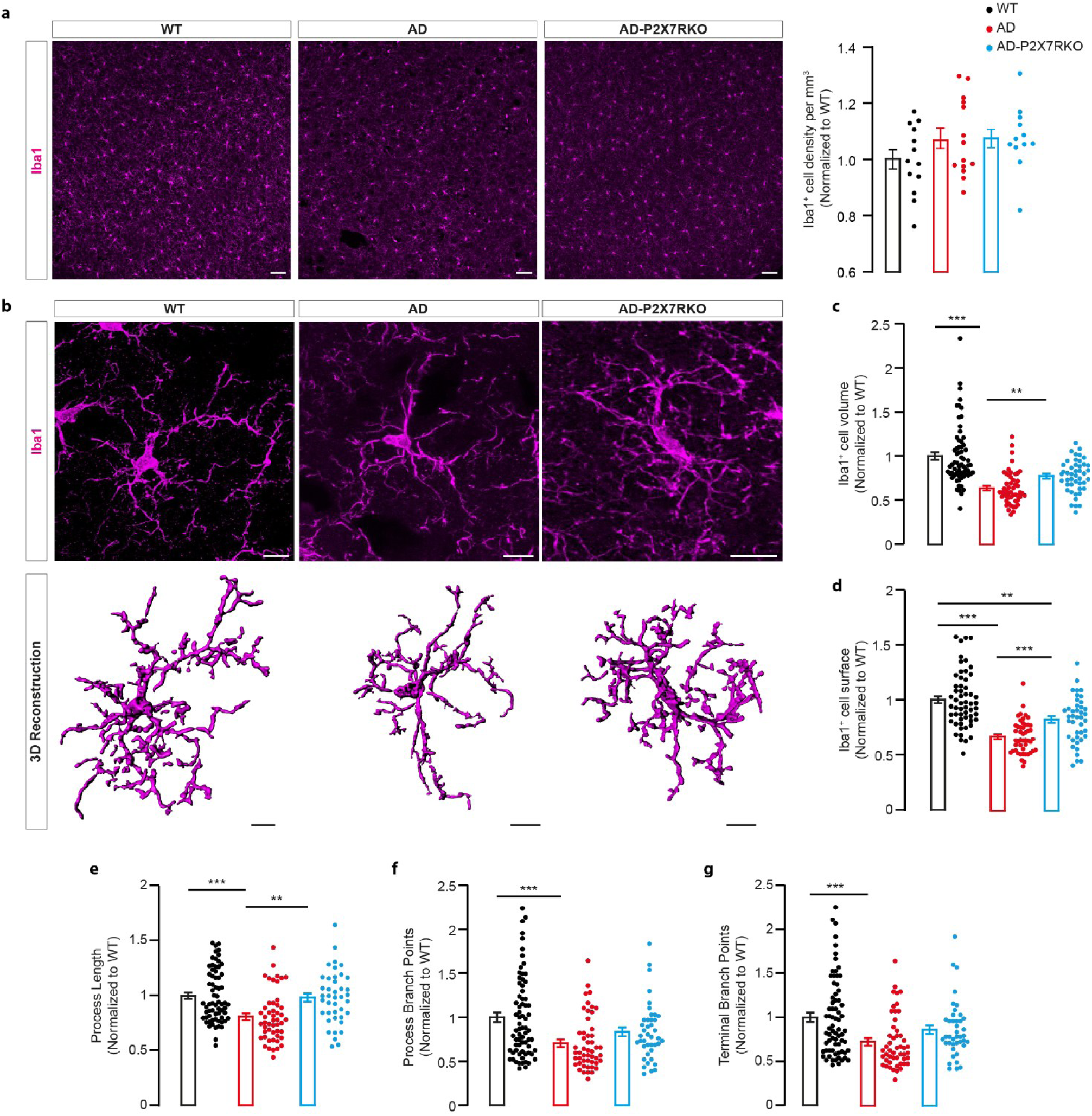
P2X7R drives early microglia morphological changes in SSCx L4 of 2-month-old AD mice. **a**, Immunofluorescence detection of Iba1⁺ microglia of WT (left), AD (middle), and AD-P2X7RKO (right) mice. Left, representative confocal images acquired at low magnification for microglia density assessment (scale bar, 50 µm); right, quantification of Iba1^+^ cell density in WT [n = 13 fields of view (FoVs)], AD (n = 14 FoVs), and AD-P2X7RKO (n = 12 FoVs) mice. N ≥ 3 animals. **b**, Immunofluorescence detection of Iba1⁺ microglia, as in panel a, imaged at higher magnification for morphological analysis. Top, representative confocal images (scale bar, 10 µm); bottom, three-dimensional (3D) reconstructions (scale bar, 10 µm). **c–g**, Quantification of Iba1⁺ microglia volume (c; WT, n = 64 cells; AD, n = 51; AD-P2X7RKO, n = 43), surface area (d; WT, n = 57; AD, n = 48; AD-P2X7RKO, n = 43; one-way ANOVA, Dunnett’s multiple comparisons test), process length (e; WT, n = 66; AD, n = 50; AD-P2X7RKO, n = 41), branch points (f; WT, n = 71; AD, n = 50; AD-P2X7RKO, n = 41), and terminal branch points (g; WT, n = 71; AD, n = 50; AD-P2X7RKO, n = 41). N ≥ 3 mice/genotype. Data are presented as mean ± s.e.m. normalized to WT; statistical analysis was performed using Kruskal–Wallis test followed by Dunn’s post hoc test. For panel a, one-way ANOVA followed by Dunnett’s multiple comparisons test was applied. \*\**P* < 0.01; \*\*\**P* < 0.001.

To resolve early structural changes, we performed high-resolution Imaris reconstructions of Iba1⁺ microglia in L4 of the SSCx (Fig. 2b). Strikingly, before Aβ deposition, AD microglia show reduced cell volume and surface area (Fig. 2b,c) and shorter processes (Fig. 2d), consistent with arbor simplification and an amoeboid-like phenotype. These changes are largely rescued in AD-P2X7RKO mice. Branch points (Fig. 2e) and terminal tips (Fig. 2f) are also reduced in AD compared with WT and partially restored by P2X7R deletion. At 6 months, microglia alterations persist in AD mice, with only partial improvement in AD-P2X7RKO animals (Extended Data Fig. 2b–f), again suggesting reduced efficacy of genetic P2X7R deletion once Aβ burden has accumulated.

### P2X7R activation drives aberrant synaptic engulfment via complement-dependent microglial phagocytosis

Microglia refine neural circuits by engulfing excess or dysfunctional synapses, a process essential for brain development and plasticity. Under pathological conditions, however, this function becomes aberrant and contributes to synaptic pathology^22^. In AD patients, synaptic imbalance arises early and correlates with cognitive decline^23^, yet the upstream signals driving synaptic dysfunction remain poorly defined.

We hypothesize that the eATP–P2X7R signaling axis, by promoting early microglia activation, alters microglia-mediated synaptic remodeling. To test this, we first quantified CD68⁺ structures within Iba1⁺ cells in L4 of SSCx, as a marker of microglial phagocytic activity^24^. At 2 months, the number of microglial CD68⁺ structures is significantly increased in AD compared with controls; importantly, it is restored to control levels in AD-P2X7RKO animals (Fig. 3a). This phenotype becomes even more pronounced at 6 months and remains P2X7R-dependent (Extended Data Fig. 3a). These data indicate an early and sustained enhancement of microglial phagocytic activity in AD mice, which critically depends on P2X7R signaling.

**Figure 3.**
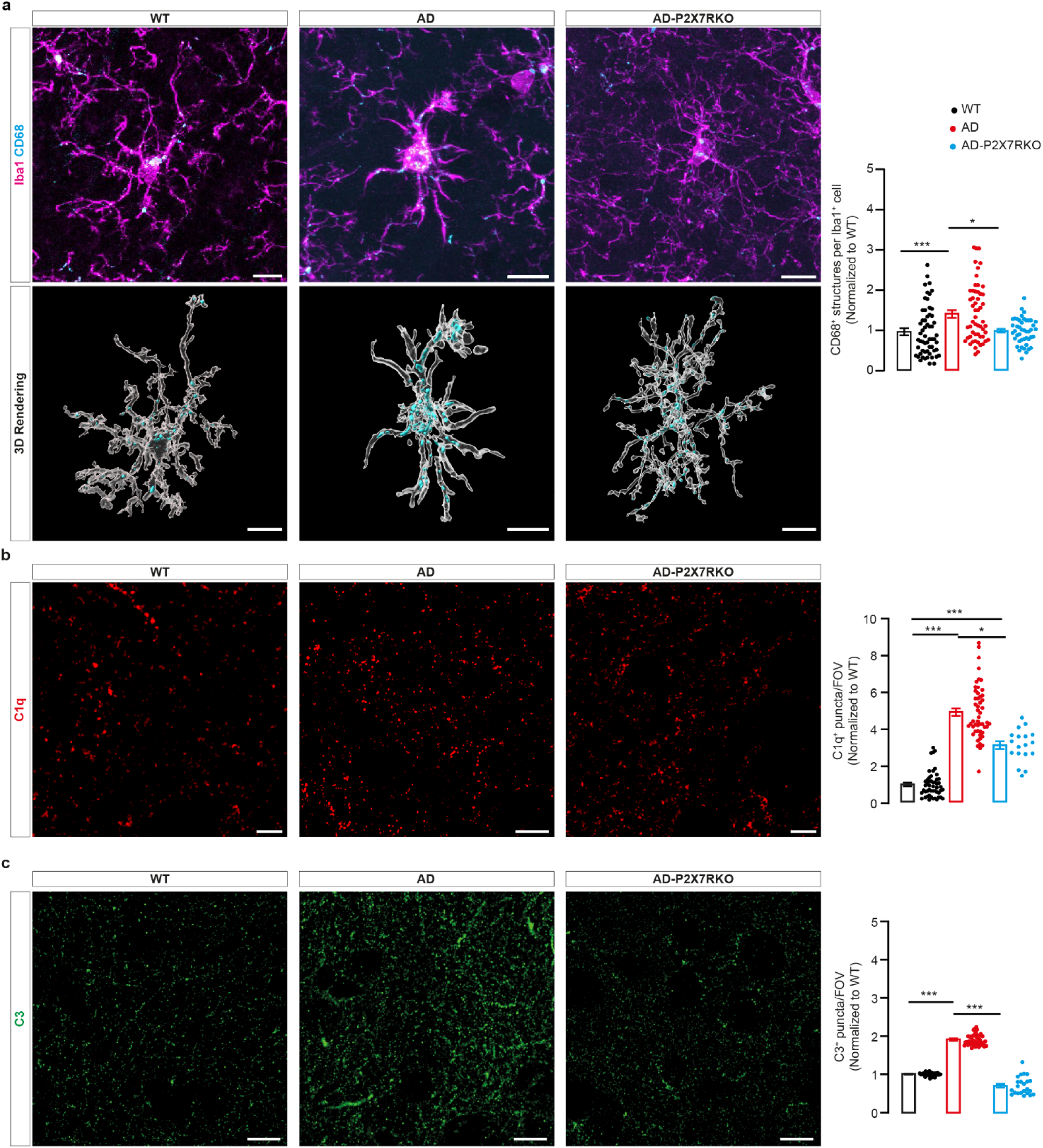
P2X7R drives early microglia phagocytic activity and complement activation in SSCx L4 of 2-month-old AD mice. **a**, Immunofluorescence detection of CD68⁺ structures within Iba1⁺ microglia of WT (left), AD (middle), and AD-P2X7RKO (right) mice. Top, representative confocal images of microglia immunolabeled for CD68 (cyan) and Iba1 (magenta) (scale bar, 10 µm); bottom, 3D reconstructions (scale bar, 10 µm); right, quantification of CD68⁺ structures per Iba1⁺ cell in WT (n = 58), AD (n = 55), and AD-P2X7RKO (n = 45) mice. N ≥ 3 mice/genotype. **b,** Immunofluorescence detection of C1q⁺ puncta in WT (left), AD (middle), and AD-P2X7RKO (right) mice. Left, representative confocal images (scale bar, 10 µm); right, quantification of C1q^+^ puncta per FoV in WT (n = 50 FoVs), AD (n = 51 FoVs), and AD-P2X7RKO (n = 18 FoVs) mice. N ≥ 3 mice/genotype. **c,** Immunofluorescence detection of C3^+^ puncta in WT (left), AD (middle), and AD-P2X7RKO (right) mice. Left, representative confocal images (scale bar, 10 µm); right, quantification of C3^+^ puncta per FoV in WT (n = 29 FoVs), AD (n = 45 FoVs), and AD-P2X7RKO (n = 23 FoVs) mice. N ≥ 3 mice/genotype. Data are presented as mean ± s.e.m. normalized to WT; statistical analysis was performed using Kruskal–Wallis test followed by Dunn’s post hoc test. *P < 0.05; ***P < 0.001.

Synaptic pruning by microglia is regulated by the complement proteins C1q and C3, which act as “eat-me” signals on vulnerable synapses^25^. Consistent with enhanced phagocytosis, AD mice show a marked increase in C1q⁺ and C3⁺ puncta at 2 months, and both values are significantly reduced in AD-P2X7RKO mice (Fig. 3b,c), with similar results at 6 months (Extended Data Fig. 3b,c). To investigate whether inhibitory or excitatory synapses were selectively targeted for microglia phagocytosis, we analyzed the engulfment of specific synaptic markers by Iba1^+^ cells. In AD mice, gephyrin⁺ inhibitory postsynaptic compartments are increasingly internalized within CD68⁺ microglia at both 2 and 6 months, a phenotype partially rescued by P2X7R deletion (Fig. 4a,b; Extended Data Fig. 4a,b). This is accompanied by a reduction in overall gephyrin⁺ puncta in the AD SSCx, a phenotype mitigated by P2X7R deletion (Fig. 4c; Extended Data Fig. 4c). Gephyrin–C1q co-localization is increased in 2-month-old AD mice, indicating complement-mediated tagging of inhibitory synapses, and is again blunted by P2X7R deletion (Fig. 4d). Similarly, gephyrin–C1q co-localization is increased in 6-month-old AD mice, and the P2X7R deletion partially rescues this phenotype (Extended Data Fig. 4d).

**Figure 4.**
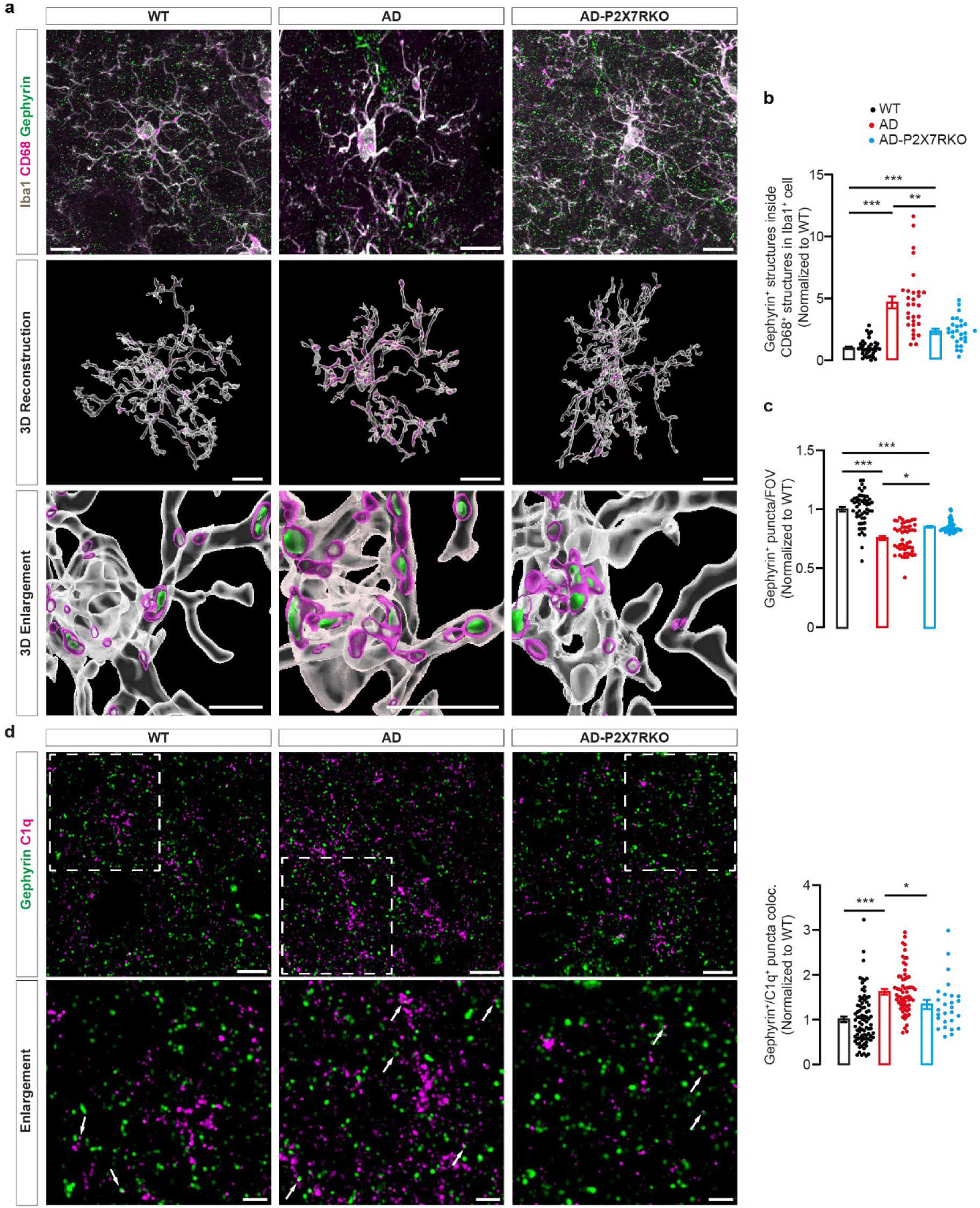
P2X7R drives early complement-mediated engulfment of inhibitory synapses by microglia in SSCx L4 of 2-month-old AD mice. **a,** Immunofluorescence detection of CD68⁺ structures colocalized with gephyrin^+^ synapses within Iba1⁺ microglia in WT (left), AD (middle), and AD-P2X7RKO (right) mice. Top, representative confocal images of Iba1⁺ (grey) microglia also immunolabeled for CD68 (magenta) and gephyrin (green) (scale bar, 10 µm); middle, 3D reconstructions (scale bar, 10 µm); bottom, 3D reconstruction enlargement (scale bar, 5 µm). **b**, Quantification of CD68⁺ structures colocalized with gephyrin^+^ synapses per Iba1⁺ cell in WT (n = 36 FoVs), AD (n = 30 FoVs), and AD-P2X7RKO (n = 27 FoVs) mice. N ≥ 3 mice/genotype. **c**, Quantification of gephyrin^+^ puncta per FoV in WT (n = 50 FoVs), AD (n = 50 FoVs), and AD-P2X7RKO (n = 50 FoVs) mice. N ≥ 3 mice/genotype. **d**, Immunofluorescence detection of C1q⁺ puncta colocalized with gephyrin⁺ synapses in WT (left), AD (middle), and AD-P2X7RKO (right) mice. Top left, representative confocal images of C1q⁺ (magenta) and gephyrin⁺ (green) puncta (scale bar, 5 µm), with at the bottom the enlargement on the dashed box, white arrows indicate representative colocalizations (scale bar, 2 µm); right, quantification of C1q⁺ puncta colocalized with gephyrin⁺ ones in WT (n = 84 FoVs), AD (n = 65 FoVs), and AD-P2X7RKO (n = 26 FoVs) mice. N ≥ 3 mice/genotype. Data are presented as mean ± s.e.m. normalized to WT; statistical analysis was performed using Kruskal–Wallis test followed by Dunn’s post hoc test. \**P* < 0.05; \*\**P* < 0.01; \*\*\**P* < 0.001.

Excitatory synapses are similarly affected. PSD95⁺ puncta are more frequently engulfed in CD68⁺/Iba1⁺ cells in AD mice compared with controls, at both ages. This aberrant phagocytosis is fully reversed by P2X7R deletion at 2 months (Fig. 5a,b), but not at 6 months (Extended Data Fig. 5a,b). Accordingly, PSD95⁺ density is reduced in AD brains with a partial recovery in AD-P2X7RKO animals at both 2 (Fig. 5c) and 6 (Extended Data Fig. 5c) months. PSD95–C1q co-localization follows the same pattern as gephyrin, increasing in AD mice and being restored to control levels upon P2X7R deletion (Fig. 5d; Extended Data Fig. 5d).

**Figure 5.**
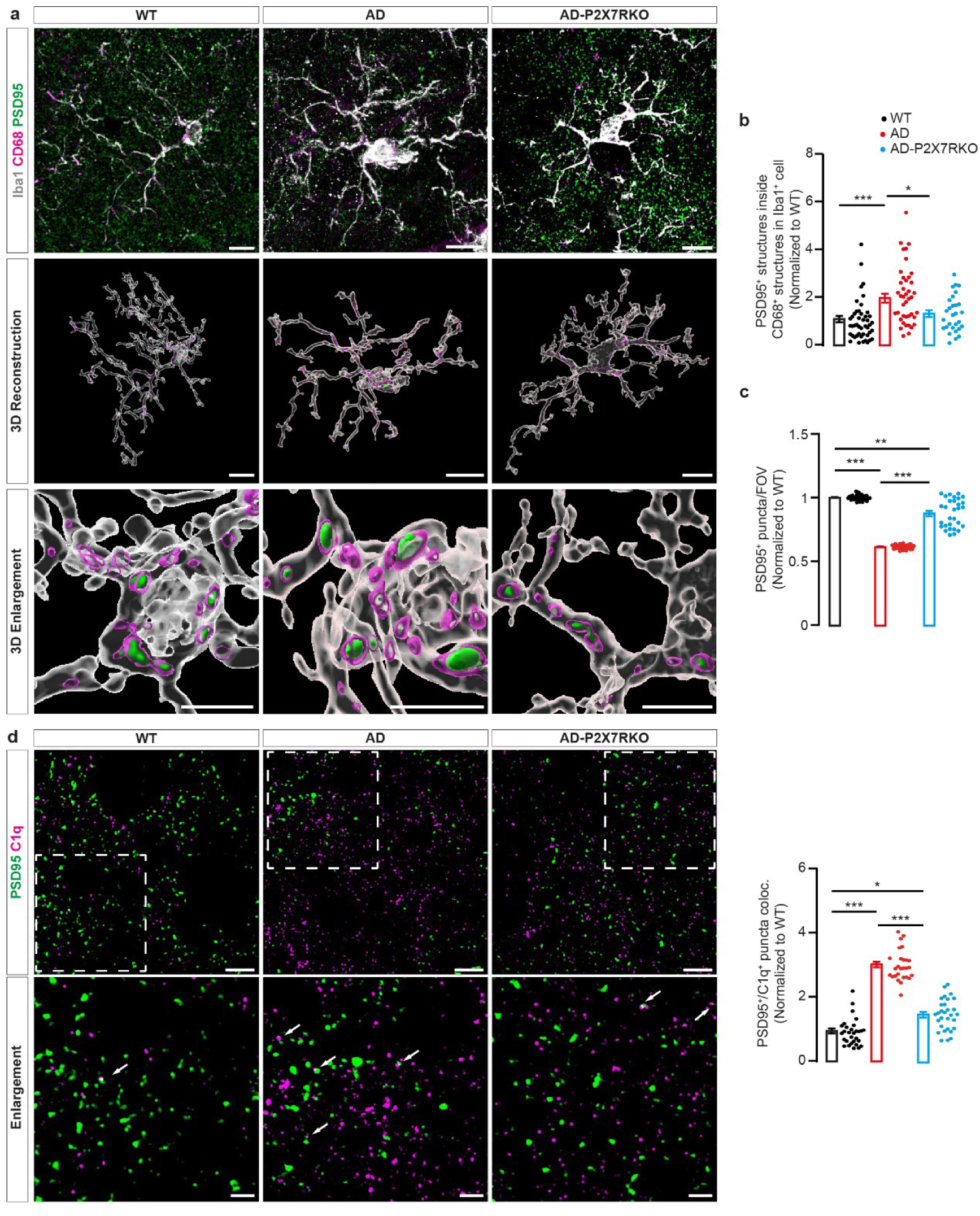
P2X7R drives early complement-mediated pruning of excitatory synapses in SSCx L4 of 2-month-old AD mice. **a,** Immunofluorescence detection of CD68⁺ structures colocalized with PSD95^+^ synapses within Iba1⁺ microglia in WT (left), AD (middle), and AD-P2X7RKO (right) mice. Top, representative confocal images of Iba1⁺ (grey) microglia also immunolabeled for CD68 (magenta) and PSD95 (green) (scale bar, 10 µm); middle, 3D reconstructions (scale bar, 10 µm); bottom, 3D reconstruction enlargement (scale bar, 5 µm). **b**, Quantification of CD68⁺ structures colocalized with PSD95^+^ puncta per Iba1⁺ cell in WT (n = 46 FoVs), AD (n = 41 FoVs), and AD-P2X7RKO (n = 29 FoVs) mice. N ≥ 3 mice/genotype. **c**, Quantification of PSD95^+^ puncta per FoV in WT (n = 33 FoVs), AD (n = 29 FoVs), and AD-P2X7RKO (n = 30 FoVs) mice. N ≥ 3 mice/genotype. **d**, Immunofluorescence detection of C1q⁺ puncta colocalized with PSD95⁺ synapses in WT (left), AD (middle), and AD-P2X7RKO (right) mice. Top left, representative confocal images of C1q⁺ (magenta) and PSD95⁺ (green) puncta (scale bar, 5 µm), with at the bottom the enlargement on the dashed box, white arrows indicate representative colocalizations (scale bar, 2 µm); right, quantification of C1q⁺ puncta colocalized with PSD95⁺ ones in WT (n = 30 FoVs), AD (n = 30 FoVs), and AD-P2X7RKO (n = 30 FoVs) mice. N ≥ 3 mice/genotype. Data are presented as mean ± s.e.m. normalized to WT; statistical analysis was performed using Kruskal–Wallis test followed by Dunn’s post hoc test. \**P* < 0.05; \*\**P* < 0.01; \*\*\**P* < 0.001.

### P2X7R-dependent microglia activation disrupts PNNs in AD mice

Beyond synaptic pruning, microglia also remodel the ECM, which shapes synaptic architecture and regulates neuronal excitability. PNNs — specialized ECM lattices surrounding mainly parvalbumin-positive (PV⁺) interneurons — stabilize inhibitory circuits and restrict plasticity^26^. Notably, PNN alterations have been reported in AD patients and animal models^27^, but the mechanisms underlying their early disruption remain poorly understood.

To test whether early microglia reactivity via P2X7R contributes to PNN alterations, we examined PNNs in L4 of the SSCx in WT, AD, and AD-P2X7RKO mice. Wisteria floribunda agglutinin (WFA) staining reveals a reduction in PNN density in 2-month-old AD mice compared with WT, a phenotype that persists at 6 months and is partially rescued in AD-P2X7RKO animals (Fig. 6a; Extended Data Fig. 6a).

**Figure 6.**
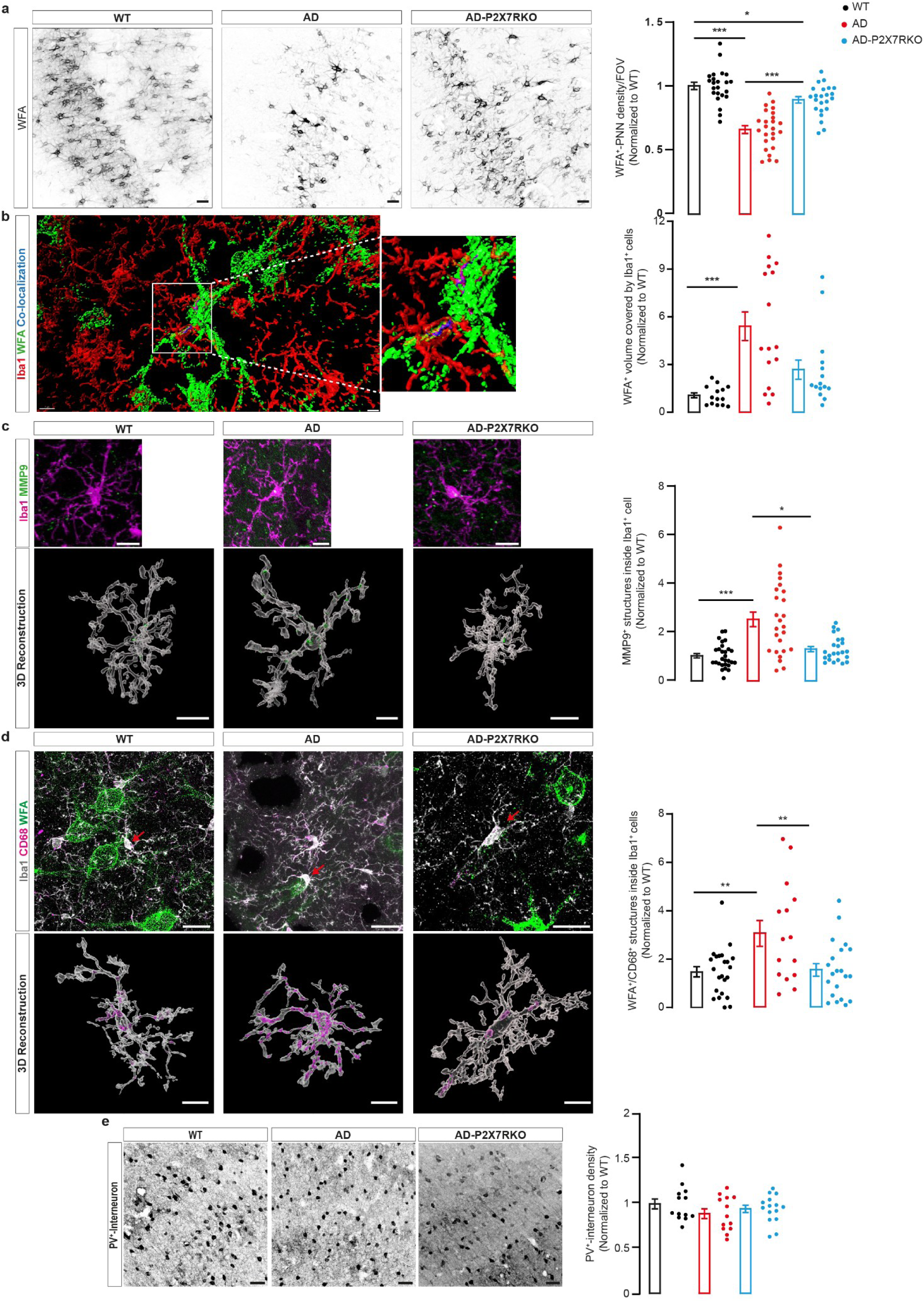
Microglia-mediated degradation of PNNs in SSCx L4 of 2-month-old AD mice depends on P2X7R. **a,** Left, WFA⁺-PNN structures in WT (left), AD (middle), and AD-P2X7RKO (right) mice. Representative confocal images (scale bar, 50 µm). Right, quantification of WFA⁺ structures per FoV in WT (n = 23 FoVs), AD (n = 25 FoVs), and AD-P2X7RKO (n = 23 FoVs) mice. N ≥ 3 mice/genotype. **b,** Left, representative 3D reconstruction of Iba1⁺ (red) microglia and WFA⁺ structures (green, scale bar, 7 µm). Right, quantification of WFA⁺-PNNs colocalized with Iba1⁺ cells in WT (n = 14 FoVs), AD (n = 16 FoVs), and AD-P2X7RKO (n = 15 FoVs) mice. N ≥ 3 mice/genotype. **c,** Left, immunofluorescence detection of MMP9⁺ structures within Iba1⁺ microglia in WT (left), AD (middle), and AD-P2X7RKO (right) mice. Top, representative confocal images of Iba1⁺ (magenta) microglia and MMP9 (green, scale bar, 10 µm); Bottom, 3D reconstructions (scale bar, 10 µm). Right, quantification of MMP9⁺ structures within Iba1⁺ cells in WT (n = 28 FoVs), AD (n = 25 FoVs), and AD-P2X7RKO (n = 23 FoVs) mice. N ≥ 3 mice/genotype. **d,** Left, immunofluorescence detection of CD68⁺ structures colocalized with WFA⁺-PNN structures within Iba1⁺ microglia in WT (left), AD (middle), and AD-P2X7RKO (right) mice. Top, representative confocal images of Iba1⁺ (magenta) microglia also labeled for CD68 (cyan) and WFA⁺ structures (green, scale bar, 20 µm). Red arrows indicate the reconstructed microglial cell; bottom, 3D reconstructions (scale bar, 10 µm). Right, quantification of WFA⁺-PNN fragments within CD68⁺ structures in Iba1⁺ cells of WT (n = 23 FoVs), AD (n = 15 FoVs), and AD-P2X7RKO (n = 21 FoVs) mice. N ≥ 3 mice/genotype. **e,** Left, representative confocal images of PV⁺ interneurons in WT (left), AD (middle), and AD-P2X7RKO (right) mice (scale bar, 50 µm). Right, quantification of PV⁺ interneuron density per FoV in WT (n = 13 FoVs), AD (n = 13 FoVs), and AD-P2X7RKO (n = 15 FoVs) mice. N ≥ 3 mice/genotype. Data are presented as mean ± s.e.m. normalized to WT; statistical analysis was performed using Kruskal–Wallis test followed by Dunn’s post hoc. In panels a, d and e one-way ANOVA followed by Dunnett’s multiple comparisons test was applied. \**P* < 0.05; \*\**P* < 0.01; \*\*\**P* < 0.001.

Because PNN degradation likely requires direct interaction with reactive microglia, we quantified the proximity between Iba1⁺ cells and WFA⁺ structures. At 2 months, AD microglia cover a greater percentage of PNNs than WT cells, a parameter significantly reduced in AD-P2X7RKO mice (Fig. 6b). At 6 months, microglia–PNN proximity remains elevated in AD mice but is no longer reversed by P2X7R deletion (Extended Data Fig. 6b).

Mechanistically, microglia-mediated ECM remodeling involves enzymatic PNN cleavage followed by phagocytic clearance of PNN fragments. Among ECM-modifying enzymes, matrix metalloproteinase 9 (MMP9) has been specifically implicated in PNN degradation^28^ and is upregulated in reactive microglia in several neuroinflammatory conditions, including AD^29,30^. Interestingly, in 2-month-old AD mice, but not in AD-P2X7RKO animals, microglia contain a higher number of MMP9⁺ vesicles compared with WT (Fig. 6c). At 6 months, MMP9⁺ vesicles remain elevated in AD mice and this increase is attenuated in AD-P2X7RKO animals (Extended Data Fig. 6c). In parallel, CD68⁺ lysosomes in AD microglia contain more WFA⁺ fragments than in controls, indicating enhanced phagocytosis of PNN components in AD mice. This feature is absent in 2-month-old AD-P2X7RKO mice (Fig. 6d) and only partially maintained at 6 months (Extended Data Fig. 6d). A consequence of this microglia-dependent ECM remodeling is a change in the percentage of PV⁺ cells ensheathed by PNNs in 2-month-old AD mice (Extended Data Fig. 6e), although PV⁺ interneuron density remains unchanged (Fig. 6e). By 6 months, however, the PV⁺/PNN⁺ ratio is re-established across genotypes, but the total number of PV⁺ cells is reduced in AD mice (Extended Data Fig. 6e,f). PV^+^ interneuron loss is partially rescued by P2X7R deletion (Extended Data Fig. 6f), suggesting that early eATP–P2X7R signaling drives PNN loss around PV⁺ interneurons, rendering them more vulnerable at later disease stages.

### P2X7R signaling drives early cortical hyperexcitability in AD mice

Neuronal hyperexcitability is increasingly recognized as an early sign of AD, preceding overt neurodegeneration and contributing to circuit instability. This altered network activity likely reflects a disrupted excitation–inhibition balance, arising from synaptic loss, PV⁺ interneuron dysfunction, ECM remodeling, and glial reactivity^31^.

As shown above, P2X7R-dependent microglial synaptic pruning and ECM degradation are already present in 2-month-old AD mice (see Fig. 3–6). To test whether these changes affect neuronal function, we assessed cortical excitability by 2P Ca²⁺ imaging in acute SSCx brain slices from 2-month-old mice expressing the Ca^2+^ indicator GCaMP6f. Because spontaneous activity in our brain slice preparation was low, we delivered electrical stimulation at the L4–L5 border and monitored evoked Ca²⁺ responses in L2/3 (Fig. 7a). The protocol comprised three steps: (1) low-frequency stimulation (LFS) at 0.03 Hz for a total imaging time of 2 minutes (min) (LFS1 series); (2) high-frequency stimulation (HFS) at 100 Hz followed by LFS, for a total imaging time of 2 min; and (3) LFS at 0.03 Hz delivered 10 min after HFS (LFS2 series), for a total imaging time of 2 min (Fig. 7a–c).

**Figure 7.**
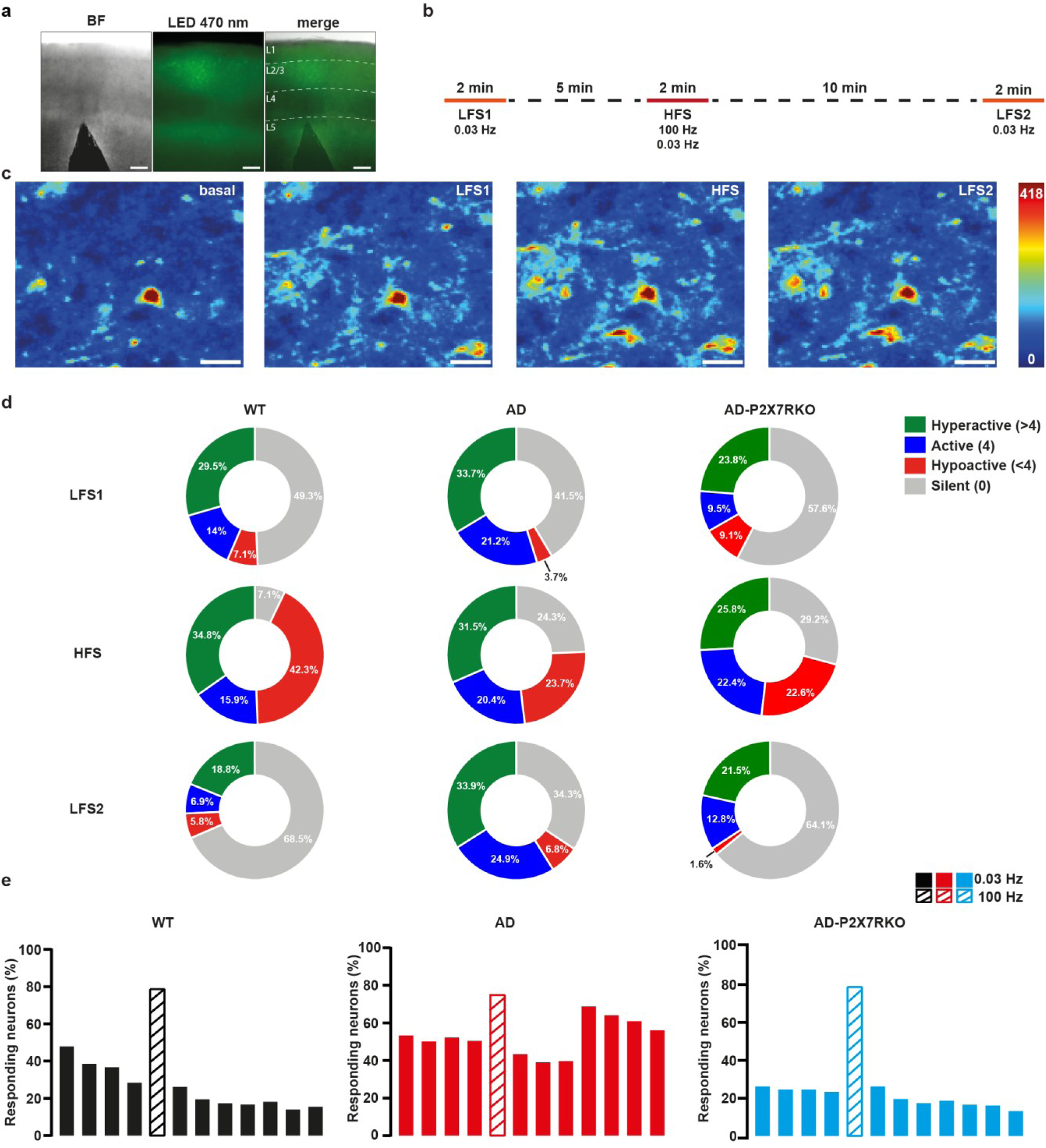
Early cortical hyperexcitability in the SSCx L2/3 of 2-month-old AD mice depends on P2X7R. **a**, Placement of the stimulation electrode and fluorescence signal. Left, representative bright-field (BF) image showing electrode placement at the L4/L5 border of SSCx. Middle, representative fluorescence image (LED 470) of GCaMP6f signal in SSCx L2/3. Right, merged image with indicated the different cortical layers (L1–5). **b,** Scheme of the imaging protocol coupled to electrical stimulation. Solid lines indicate recording periods; dotted lines indicate recovery times between imaging periods. **c,** Representative pseudo-color two-photon maximum Z-projections (20 frames) of the GCaMP6f signal from an AD SSCx mouse slice, showing basal fluorescence and responses upon low-frequency stimulation series 1 (LFS1), high-frequency stimulation series (HFS) and low-frequency stimulation series 2 (LFS2). Scale bar, 20 µm. **d,** Pie charts showing the percentage of responsive neurons classified as hyperactive (>4 Ca²⁺ events, green), active (4 Ca²⁺ events, blue), hypoactive (<4 Ca²⁺ events, red), or silent (0 Ca²⁺ events, grey) in WT, AD, and AD-P2X7RKO mice. Top row, LFS1; middle row, HFS; bottom row, LFS2. Data from WT (n = 311 neurons), AD (n = 513), and AD-P2X7RKO (n = 238). N ≥ 4 mice/genotype. Statistical comparisons: • LFS1: WT vs AD ** *P* < 0.01; WT vs AD-P2X7RKO * *P* < 0.05; AD vs AD-P2X7RKO *** *P* < 0.001. • HFS: WT vs AD *** *P* < 0.001; WT vs AD-P2X7RKO *** *P* < 0.001; AD vs AD-P2X7RKO * *P* < 0.05. • LFS2: WT vs AD *** *P* < 0.001; WT vs AD-P2X7RKO * *P* < 0.05; AD vs AD-P2X7RKO *** *P* < 0.001. χ² test followed by pairwise comparisons with Bonferroni correction. **e,** Bar histograms showing the percentage of responsive neurons to each stimulation (LFS or HFS) in WT, AD, and AD-P2X7RKO mice. Data from WT (n = 311 neurons), AD (n = 513), and AD-P2X7RKO (n = 238). N ≥ 4 mice/genotype. Statistical comparisons: WT vs AD ** P < 0.001; WT vs AD-P2X7RKO, P = 0.05; AD vs AD-P2X7RKO *** *P* < 0.001. Kolmogorov–Smirnov test followed by Bonferroni correction, α = 0.0167.

Neurons were categorized based on their activity profile for each series: hyperactive (>4 Ca²⁺ events), active (4 Ca²⁺ events), hypoactive (<4 Ca²⁺ events), or silent (no Ca^2+^ transients during that series, but active in at least one of the others). In AD slices, the proportion of hyperactive and active neurons during both LFS1 and LFS2 is higher than in WT, indicating heightened basal and post-stimulation excitability (Fig. 7d). In WT slices, HFS recruits 80% of neurons but is followed by a sharp reduction during LFS2, with only ∼20% of neurons remaining active, consistent with adaptive downscaling^32^ (Fig. 7d,e). In contrast, AD slices show a paradoxical increase in responsive neurons during LFS2, revealing sustained hyperexcitability (Fig. 7e).

To determine whether this phenotype depends on P2X7R, we analyzed SSCx brain slices from AD-P2X7RKO mice. During LFS1, fewer neurons respond compared with both WT and AD (Fig. 7d,e), suggesting that P2X7R contributes to basal neuronal excitability. Nevertheless, following HFS, neuronal recruitment (∼80%) is comparable to both WT and AD mice (Fig. 7e). Crucially, during LFS2, AD-P2X7RKO slices mirror WT responses, with a decline in active neurons, reflecting normal adaptation (Fig. 7e). Thus, P2X7R deletion prevents the persistent hyperexcitable state observed in AD mice.

## Discussion

The prodromal phase of AD is increasingly recognized as a dynamic time-window during which subtle, yet decisive, molecular, cellular, and network changes emerge before overt Aβ plaque deposition or neurodegeneration. Neuroinflammation is a recurring feature of AD, yet whether it represents a downstream reaction to amyloid pathology or an upstream driver of both synaptic and circuit dysfunction remains poorly understood^33^. Among candidate mediators, P2X7R has been repeatedly implicated in brain inflammation. It is upregulated in microglia of multiple AD models and in *postmortem* AD brains, particularly around Aβ plaques^12,13^, where it promotes cytokine release, ROS production, and glia cell migration toward amyloid plaques. Genetic studies also support a modulatory role of P2X7R in AD, with distinct single-nucleotide polymorphisms (SNPs) associated with either anti-inflammatory responses or enhanced clearance of Aβ. Furthermore, genetic and pharmacological blocking of P2X7R reduces amyloid pathology while improving synaptic plasticity and cognitive performance. However, the timing and functional impact of P2X7R activation in AD remain unclear^14^.

Here, we identify the eATP–P2X7R axis as a causal and upstream mechanism linking early microglia activation to synaptic pathology and neuronal hyperexcitability in the PS2APP AD mouse model. Direct *in vivo* measurements show that cortical eATP accumulation is detectable by 2 months of age — before Aβ deposition and behavioral decline — and persists throughout disease progression. This provides the first evidence that chronic eATP elevation extends beyond acute brain injury^34^ and is a defining feature of neurodegenerative conditions. This early eATP surge appears independent of transcriptional changes in P2X7R or ectonucleotidases, yet critically depends on the presence of P2X7R. Indeed, P2X7R deletion in AD mice normalizes eATP levels, supporting the existence of a feed-forward loop in which the receptor both senses and sustains high eATP, either through its pore-forming activity or indirectly via other channels^11,35^.

Mechanistically, eATP–P2X7R activation triggers a stereotyped inflammasome response, with NLRP3 upregulation and increased production of IL-1β, TNF-α, and IL-6 in the brain of AD mice at 2 months of age. By 6 months, TNF-α and IL-6 levels return to control values, despite ongoing IL-1β elevation, consistent with previous reports indicating that IL-1β acts as an early and persistent driver of AD-related neuroinflammation. IL-1β can amplify amyloid precursor protein (APP) processing, tau phosphorylation, and excitotoxicity, thereby sustaining a chronic pro-inflammatory *milieu* even when other inflammatory mediators normalize. In contrast, TNF-α and IL-6 display more transient or context-dependent dynamics during AD progression^36^. Importantly, these distinct cytokine profiles may have implications for staging disease-modifying interventions. Strikingly, P2X7R deletion normalizes cytokine levels and NLRP3 protein expression at both ages, underscoring its role as a gatekeeper of immune activation in early AD.

Activation of P2X7R and cytokine release accompany microglia reactivity^37^. In the AD brain, morphological alterations of microglia are typically associated with Aβ plaque appearance^33^. However, single-cell transcriptomic studies show that microglia lose their homeostatic identity and adopt disease-associated states early in AD^38,39^. Our data provide a mechanistic explanation: eATP functions as a DAMP that triggering innate immune responses via P2X7R activation in the absence of an overt amyloid phenotype. Indeed, before Aβ plaque accumulation, AD microglia exhibit reduced area, volume, and process complexity — features consistent with an amoeboid-like, surveillance-deficient state. P2X7R deletion partially rescues this phenotype, with full recovery of cell area and volume, indicating that P2X7R engagement is sufficient to initiate microglia transformation. Other pathways, however — such as activation of additional purinergic receptors (P2Y12, P2X4)^40^, neurotransmitter imbalances^41^, release of other neuron-derived DAMPs (HMGB1, CX3CL1)^42,43^, epigenetic priming by inflammatory cues^44^, or further ECM degradation^28^ — may act in parallel to sustain microglia reactivity once established.

Functionally, we find that AD reactive microglia show an increase in CD68⁺ phagolysosomal structures, fully rescued by P2X7R deletion at both early and late AD stages, indicating that early purinergic stress directly primes microglia for phagocytosis. The consequences of this activity, however, may depend on context: while under physiological conditions it promotes clearance of cellular debris or pruning of excessive synapses, in AD the combination of Aβ and chronically elevated eATP biases microglia toward maladaptive, P2X7R-dependent phagocytosis, with exaggerated synaptic elimination and ECM disruption^33^.

Aberrant synaptic engulfment is a common hallmark of neurodevelopmental and neurodegenerative disorders^45^, with the complement cascade playing a crucial role in the process^46^. In pathological conditions, both C1q and C3 mediate dysfunctional synapse elimination: C1q drives the removal of inhibitory synapses during aging^47^ and contributes to both excitatory and inhibitory loss in AD^48,49^, whereas C3 mediates excitatory pruning in multiple sclerosis^50^. Consistent with this, we find that both C1q and C3 increase in AD brains, with consequent potentiated engulfment of both gephyrin⁺ inhibitory and PSD95⁺ excitatory synapses by microglia. This phenotype is rescued in AD-P2X7RKO mice, placing eATP–P2X7R upstream of complement-mediated synapse elimination. Interestingly, we observe a differential recovery of C1q (partial) versus C3 (full) levels upon P2X7R deletion, suggesting distinct regulatory mechanisms. While C1q is produced by multiple brain cell types and is responsive to oligomeric Aβ as well as other DAMPs^51^, C3 is mainly produced by astrocytes and largely induced by pro-inflammatory cytokines^52^. Thus, the full rescue of C3 levels likely reflects the robust suppression of cytokine-driven inflammation observed early in AD-P2X7RKO animals. Importantly, inhibitory and excitatory synapses show divergent sensitivity to purinergic stress. Inhibitory gephyrin⁺ synapses remain only partially protected by P2X7R deletion, suggesting that their clearance may proceed via alternative mechanisms, such as neurotransmitter-driven pathways^53^. In contrast, excitatory PSD95⁺ synapses are fully rescued by P2X7R deletion despite residual C1q deposition, indicating a stricter dependence on P2X7R signaling.

Besides synaptic remodeling, microglia can also directly engulf ECM components^3^. PNNs, which stabilize PV⁺ interneurons and thereby preserve excitatory–inhibitory balance^27^, are disrupted in both AD models and patients^54^, together with PV⁺ interneuron dysfunction^55^.

Notably, we observe high microglia phagocytic activity toward PNNs already in 2-month-old AD mice, together with an increase in the number of MMP9⁺ vesicles within microglia. This aligns with the established role of MMP9 and phagocytosis in PNN degradation^28^ and is consistent with previous reports showing MMP9 upregulation in the AD brain^29,30^. Of note, P2X7R deletion fully restores these phenotypes at 2 months, but only partially at 6 months, suggesting that purinergic signaling is critical in early phases, whereas later disease stages are sustained by additional detrimental factors, including Aβ plaque–driven inflammation.

PNN loss is accompanied by a reduction in PV⁺ interneurons, which becomes evident at 6 months, although the process of PNN degradation began as early as 2 months. This effect is partially rescued by P2X7R deletion. These findings are consistent with growing evidence that PNN degradation destabilizes PV⁺ interneuron function and contributes to impaired inhibitory tone in both AD and related neuropsychiatric conditions^56^.

Together, aberrant synaptic pruning and PNN loss compromise inhibitory control of cortical circuits, providing a structural basis for altered brain excitability. Interestingly, cortical hyperexcitability and subclinical seizures are increasingly recognized in prodromal AD patients and animal models^31^, and our mice likewise exhibit early hyperexcitability across multiple experimental paradigms^57–59^. Altogether our data provide a mechanistic bridge: early purinergic stress and P2X7R activation lead to maladaptive microglia pruning, PNN loss, and consequent network disinhibition, culminating in persistent hyperexcitability. 2P Ca²⁺ imaging demonstrates that these early structural alterations translate into measurable neuronal network dysfunction. In SSCx brain slices from 2-month-old mice, AD animals display a higher percentage of responsive neurons upon successive electrical stimulation. Crucially, P2X7R deletion restores adaptive downscaling of neuronal responses after repetitive stimulation, preventing the paradoxical increase in activity observed in AD mice. Of note, AD-P2X7RKO animals exhibit fewer responsive neurons compared with WT mice, suggesting that P2X7R may play an important, previously unrecognized role in regulating neuronal activity. Importantly, neuronal hyperexcitability in AD mice emerges before Aβ plaque formation, linking purinergic signaling to early circuit dysregulation. This is consistent with prior studies associating ATP release with increased synaptic activity^60^ or neuronal injury^9^.

By demonstrating that early neuronal dysfunction in AD is P2X7R-dependent and reversible, our study provides compelling evidence that purinergic signaling is not a bystander but a causal node in the early disruption of neuronal network homeostasis. Altogether, our data point to P2X7R and eATP levels as potential early biomarkers of AD pathogenesis, and recent advances in radioactive P2X7R ligands for PET imaging provide a concrete translational route for their clinical application^61,62^.

From a therapeutic perspective, our work positions P2X7R as an attractive target for early intervention in AD. Importantly, brain-penetrant P2X7R antagonists are currently in clinical development for various neuroinflammatory conditions, offering a realistic opportunity for drug repurposing^63,64^.

In conclusion, we propose that brain accumulation of eATP and the consequent P2X7R activation serve as a pivotal signaling axis linking immune activation, microglia reactivity, and circuit dysfunction in the earliest phases of AD. Targeting the eATP–P2X7R axis may therefore offer a unique opportunity to delay or prevent the transition from silent pathology to symptomatic neurodegeneration.

## Methods

### Mouse Models

All experimental procedures were conducted in accordance with the European Directive 2010/63/EU in compliance with the ARRIVE guidelines. Protocols were approved by the University of Padua Animal Care Committee and authorized by the Italian Ministry of Health (authorization numbers: 69/2022-PR, 174/2023-PR, D2784.N.AZR). Mice were housed under standard conditions: a 12-hour light–dark cycle (lights on from 7 a.m. to 7 p.m.), room temperature kept at 22 °C, and 60% relative humidity.

Homozygous PS2APP (B6.152H) transgenic mice were generously provided by L. Ozmen (F. Hoffmann-La Roche Ltd., Basel, Switzerland; MTA 06-02-22-UniPD). The B6.152H line co-expresses human βAPP751 with the Swedish double mutation (K670N/M671L) under the control of the mouse Thy1.2 promoter, and human PS2 carrying the N141I mutation under the control of prion protein (PrP) promoter.

Only female mice were used in all experiments, consistent with sex-specific phenotypes previously reported in the PS2APP model^1^, also reflecting the higher prevalence of AD in women (Alzheimer’s Association, 2015). Experimental cohorts included mice aged 2 to 9 months. Age-matched C57BL/6J wild-type (WT) mice were used as controls.

AD-P2X7RKO mice were generated by Memmia Srl by crossing the P2X7 receptor knockout (P2X7R-KO) line^2^ with B6.152H mice. The newly created AD-P2X7RKO line was evaluated in accordance with the guidelines of the Italian Ministry of Health. A minimum of seven males and seven females, sampled from more than one litter, were monitored immediately after birth, around weaning, after sexual maturity, and at six months of age (the last experimental time point of interest). Observations were carried out for at least two reproductive cycles from the F2 generation onward. The results confirmed a healthy phenotype for the animals.

### *In Vivo* Imaging of Extracellular ATP

Brain-wide expression of the eATP bioluminescent probe pmeLUC was obtained by retro-orbital injection of AAV PHP.eB-pmeLUC (titer 10^12^ vg/mL, diluted 1:3 in filtered 0.9% NaCl) under isoflurane anesthesia. Thirty days later, mice were anesthetized (4% isoflurane) and the skull was exposed. Imaging was conducted using an IVIS-100 system (Xenogen Corp., Alameda, CA, USA) equipped with a heated stage and isoflurane manifold (isoflurane maintained between 1.5-2%). Animals were intraperitoneally injected with Beetle Luciferin, potassium salt (150 mg/Kg; E1605, Promega Corp., Madison, WI, USA) and bioluminescence was recorded (300 s, repeated for a minimum of 4 times; total imaging time of minimum 20 minutes). Luminescence values were normalized to probe cDNA levels quantified by RT-PCR. Animals were sacrificed at the end of the recording session, with brains rapidly frozen in liquid nitrogen for RT-PCR analysis.

### Cytokine measurements

Cortical tissues (∼ 500 mg) were resuspended in 500 µl of saccharose saline plus 0.05% Igepal (Sigma-Aldrich, St. Louis, MO, USA) and homogenized with 30 potter hits. Homogenates were then centrifuged at 3,000 rpm for 20 min at 4 °C, and the supernatants were diluted 1:5 in sample diluent SD09 (Bio-Techne, Minneapolis, MN, USA). Detection of IL-1β, TNF-α and IL-6 protein levels was performed using the Ella^TM^ automated Immunoassay system (Bio-Techne) with the mouse Simple Plex Cartridge Kit (SPCKA-MP-005977, Bio-Techne), according to the manufacturer’s instructions.

### RNA Quantification (RT-qPCR)

Total RNA was isolated from brain cortex homogenates using 1 mL TRIzol Reagent, followed by chloroform phase separation, according to the manufacturer’s instructions (Thermo Fisher Scientific, Waltham, MA, USA). A NanoDrop™ ND-1000 UV-Vis spectrophotometer (Isogen Life Science B.V., Utrecht, Netherlands) was used to quantify RNA. cDNA was synthesized from 1 µg of RNA using the RT^2^ First Strand Kit (Qiagen, Hilden, Germany), according to the manufacturer’s instructions. qPCR analysis was performed with the Gene expression master mix (Thermo Fisher Scientific) using probes for mouse *p2rx7* (assay no. Mm00440578, Applied Biosystems, Foster City, CA, USA), *Entpd1* (assay no. Mm00515447, Applied Biosystems), *Nt5e* (assay no. Mm 00501910, Applied Biosystems) genes. Experimental reactions were conducted by preincubation (95 °C for 10 min) and amplification (95 °C for 15 s and 60 °C for 60 s) for 40 cycles. *GAPDH* gene (assay no. 4326317E, Thermo Fisher Scientific) was used for normalization. Gene expression was assessed using the StepOne Real-Time PCR system (Applied Biosystems), and relative gene expression was calculated using the comparative 2^−ΔΔCt^ method and expressed as fold change. All reactions were run in triplicate.

### Western blotting

Whole mouse brains were extracted in cold phosphate-buffered saline (PBS), and cortices were dissected and homogenized in RIPA buffer (50 mM Tris, 150 mM NaCl, 1% Triton X-100, 0.5% deoxycholic acid, 0.1% SDS, 1 mM DTT, pH 7.5) supplemented with protease and phosphatase inhibitors (F. Hoffmann-La Roche Ltd., Basel, Switzerland), using TissueLyser II (Qiagen) at 30 Hz for 2 min. Homogenates were centrifuged at 15,000 rpm for 15 min at 4 °C, and protein concentrations were determined with the BCA assay (Euroclone S.p.A., Milan, Italy). Equal amounts of protein (30 µg) were separated on 10% SDS–PAGE gels and transferred to nitrocellulose membranes. Membranes were blocked for 1 h at room temperature in 5% non-fat milk in Tris-Buffered Saline with Tween (TBS-T; 250 mM Tris, 1.37 M NaCl, 27 mM KCl, 0.5% Tween-20, pH 7.4) and incubated overnight at 4 °C with primary antibody against NLRP3 (rabbit, 1:1,000, AG-20B-0014, AdipoGen Life Sciences, Liestal, Switzerland) diluted in blocking buffer. GAPDH (mouse, 1:10,000, MAB374, Sigma-Aldrich) was used as loading control and detected after 1 h incubation at room temperature (RT). Following three 5 min washes in TBS-T, membranes were incubated with HRP-conjugated secondary antibodies (anti-rabbit 1:5000, AQ132P, Sigma-Aldrich, St. Louis, MO, USA; anti-mouse 1:3000, 1706516, Bio-Rad Laboratories, Hercules, CA, USA) for 1 h at RT. After additional washes, immunoreactive bands were visualized by enhanced chemiluminescence (ECL; WBLUC0500, Merck Millipore, Darmstadt, Germany) and quantified with Nine Alliance software (Uvitec, Cambridge, UK). Protein levels were normalized to GAPDH.

### Histology and Immunofluorescence

#### Tissue preparation

For histological analysis, mice were anesthetized with Alfaxan (80 mg/kg)/ xylazine (10 mg/kg) and transcardially perfused with ice-cold PBS followed by 4% paraformaldehyde (PFA, w/vol, in PBS, pH 7.4). Brains were post-fixed overnight (∼16 h) in 4% PFA at 4 °C. The tissues were then washed in PBS and stored at 4 °C with 0.025% (wt/vol) sodium azide (Sigma-Aldrich). For cryoprotection, the tissue was transferred to 30% (wt/vol) sucrose in PBS and incubated at 4 °C until it submerged (≥ 2 days). Brains were embedded in O.C.T. (Optimal Cutting Temperature compound, Polysciences, Warrington, PA, USA), and sectioned coronally at 40 µm (Cryostat, CM1850 UV, Leica Microsystems GmbH, Wetzlar, Germany). Free-floating sections were stored in cryoprotectant solution [150 g Sucrose + 5 g Polyvinyl-pyrrolidone (PVP-40) + 150 mL Ethylene glycol + 0.025% (wt/vol) sodium azide in 500 mL of 0.1M PBS] at –20 °C.

#### Staining

The SSCx area was defined based on the mouse brain atlas (Paxinos and Franklin’s. *The Mouse Brain in Stereotaxic Coordinates*). Before staining, slices were washed 3 times for 15 min each in PBS at RT on a shaker. Sections were permeabilized at RT in 0.5% Triton X-100 (Sigma-Aldrich), followed by 1 h RT blocking in 2% BSA (Sigma-Aldrich) and 0.5% Triton X-100 and incubated with primary antibodies for 48 h at 4 °C. The following primary antibodies were used: Iba1 (rabbit, 1:500, 019-19741, FUJIFILM Wako Chemicals, Virgina, USA), CD68 (mouse, 1:500; MCA1957, Bio-Rad Laboratories); PSD-95 (1:500; PSD95 sdAb - FluoTag-X2 - N3702-AF568-L, Synaptic Systems GmbH, Göttingen, Germany), gephyrin (mouse, 1:500; 147 011, Synaptic Systems GmbH), PV (rabbit, 1:500; 195 002, Synaptic Systems GmbH), C1q (rabbit, 1:500; ab182451, Abcam, Cambridge, UK), C3 (rabbit, 1:500, ab97462, Abcam), MMP9 (rabbit, 1:500, ab228402, Abcam). Alexa-conjugated secondaries diluted 1:500 in PBS (Invitrogen, Carlsbad, CA, USA: donkey anti-rabbit AF555; donkey anti-rabbit AF647; donkey anti-mouse AF488; goat anti-rat AF488; goat anti-rat AF647; donkey anti-goat AF488; PSD95 is already tagged with AzDye568) were applied for 2 h at RT. Sections were mounted in Fluoromount-G (Invitrogen). Negative controls omitted primary antibodies. To stain PNNs, sections were incubated with biotinylated Wisteria floribunda agglutinin (WFA, 1:500; B-1355-2, Vector Laboratories, Burlingame, CA, USA), followed by detection with streptavidin–Alexa Fluor 488 (S11223, Thermo Fisher Scientific).

#### Imaging

Confocal imaging was performed on a Leica SP5 microscope (Leica Microsystems, GmbH, Wetzlar, Germany) using a Plan-Apochromat 40x, NA:1.25 oil objective (microglia density analysis, morphology, phagocytosis analysis; PNN and PV analysis) or Plan-Apochromat 100x, NA:1.4 oil objective (complement and synaptic structures density; co-localization analysis of PSD95/gephyrin-C1q). The pinhole size was set to 1 Airy unit. Z-stack images were acquired with a resolution of 1024 × 1024 pixels. All imaging parameters were kept constant between control and experimental groups. Representative images were adjusted in contrast and brightness using ImageJ Software (NIH, USA).

#### Analysis

The confocal images were loaded in Fiji 1.52e (http://imagej.net/Fiji). Background was removed using a rolling-ball radius 35 pixels, and images were filtered with a median 3D filter (x, y, z radii = 3). Stacks were exported as .tiff, converted to .ims with the Imaris Converter, and imported into Imaris v9.2 (Bitplane). At 6 months, microglia analyses were restricted to plaque-distant regions.

#### Reconstruction of 3D-traced microglia

Images were acquired with a 40× (NA 1.25) oil-immersion objective on a Leica SP5 with 0.5 μm Z-step. After filtering/background subtraction, images were imported into Imaris v9.2. Starting points were generated with maximum diameter threshold 12 μm and seeding interval 1 μm. Disconnected fragments were excluded (smoothness 0.6 μm). Cells at image edges with incomplete reconstructions were manually excluded.

#### Quantification of CD68 volume and PSD95/gephyrin within Iba1 cells

Surface renderings were generated on microglia and CD68 Z-stacks using Imaris Surfaces (surface detail 0.8 for microglia and 0.3 μm for CD68). The surface–surface colocalization tool was used to count CD68 surfaces within microglia (entire image). Analysis of internalization of PSD95 or gephyrin in Iba1⁺ (microglia) lysosomes (CD68) was adapted from^3^. *Co-localization analyses.* Co-localization was calculated using the Fiji ComDet plugin (https://github.com/ekatrukha/ComDet). Images were filtered with a Mexican-hat (LoG) filter (approximate puncta size 3 pixels) and thresholded for puncta detection. Thresholds were chosen to maximize detected structures while removing background and were kept constant across analyses. Puncta were considered colocalized if the maximum distance between spot centers was < 3 pixels.

#### Quantification of microglia, PNN, and PV^+^ density

The spot-function plugin of Imaris 9.2.v was used to count cells in each confocal image. PV*^+^* and PNN density was estimated as the total number of structures obtained per imaged volume (mm^3^).

### Two-Photon Calcium Imaging

#### AAV injection

AAV9.Syn.GCaMP6f.WPRE.SV40 (Penn Vector Core, University of Pennsylvania, Philadelphia, PA, USA; Addgene#100837) was injected stereotactically into the right SSCx to express the cytosolic Ca²⁺ indicator GCaMP6f sparsely in neurons. Injections were performed at postnatal days P45-55 in mice under isoflurane anesthesia (4% induction, 1-1.5% maintenance). Carprofen (5 mg/kg, s.c.) was administered for analgesia. Depth of anesthesia was monitored by respiration rate, eyelid reflex, vibrissae movements, and reactions to tail/toe pinch. Briefly, the skin over the skull was disinfected with iodopovidone and a cut was performed along the sagittal line to expose the bone. Virus (diluted in 0.9% NaCl to 10¹² vg/mL) injections were performed after drilling two holes (0.5 mm diameter, separated by a distance of 1.5 mm, 750 nl of virus into each hole) into the skull over the right SSCx (1-1.5 mm posterior to bregma, 1.5 mm lateral to sagittal sinus, and 0.3-0.6 mm depth) using a pulled glass pipette in conjunction with a custom-made pressure injection system over a period of 2 to 5 min. At the end, the pipette was left in place for 5 more min and then gently withdrawn. After injections, the skin was sutured, and mice recovered under a heat lamp and returned to their home cage. Animals were carefully monitored in the following days for recovery and used for experiments from 2-3 weeks after injection.

#### Acute brain slice preparation

Coronal SSCx slices (280 μm) were obtained from mice at P60-70. Mice were anesthetized with isoflurane; brains were removed into ice-cold ACSF (in mM: 130 NaCl, 3 KCl, 0.5 CaCl₂, 2.5 MgCl₂, 1 NaH₂PO₄, 25 NaHCO₃, 15 glucose; pH 7.4, bubbled with 95% O₂ / 5% CO₂). Slices were cut with a vibratome (Leica VT1200S, Mannheim, Germany) in the same solution, then transferred to the recording ACSF (in mM: 130 NaCl, 3 KCl, 2.5 CaCl₂, 1 MgCl₂, 1 NaH₂PO₄, 25 NaHCO₃, 15 glucose; pH 7.4, 95% O₂ / 5% CO₂) and maintained at RT throughout experiments.

#### Electrical stimulation protocol

For LFS, square current pulses (duration 0.2 ms) were applied every 30 s (0.033 Hz) using a S-900 stimulus generator connected through a stimulus isolation unit to a bipolar coaxial electrode (10-15 kΩ, TM33CCINS, World Precision Instruments, Sarasota, FL, USA) placed in SSCx L5 near the L4 border. For HFS, one theta-burst stimulation (TBS) was delivered. It consisted of five pulses of 5 ms duration each given at 100 Hz, repeated 10 times with 200 ms start-to-start interval.

#### Imaging experiments

Ca^2+^ imaging was performed with a 2P microscope (Multiphoton Imaging System, Scientifica Ltd., Uckfield, UK) equipped with a pulsed IR laser (Chameleon Ultra 2, Coherent Inc., Santa Clara, CA, USA) tuned at 920 nm. Power at the sample was kept in the range of 10-15 mW to avoid photostimulation and photobleaching. Images were acquired at 6.1 Hz, for 2 min in each stimulation series (LFS1, HFS, LFS2), using a water-immersion objective (LUMPlan FI/IR 20×, 1.05 NA; Olympus Corporation, Tokyo, Japan). The FoV ranged between 300 x 300 µm and 150 x 150 µm depending on the zoom. Neuronal Ca²⁺ activity was recorded in SSCx L2/3.

#### Data Analysis

The detection of Regions Of Interest (ROIs) displaying Ca^2+^ elevations was performed in ImageJ 1.54m. ROIs were manually drawn around neuronal soma. Pixels within each ROI were averaged to obtain a single time course of fluorescence values, F(t). Analysis of Ca^2+^ signals was performed with the AstroResp code^4^. To compare relative changes in fluorescence between different cells, we expressed the Ca^2+^ signal for each ROI as ΔF/F_0_ = (Ft − F_0_)/(F_0_). F_0_ was defined as the 15th percentile of the whole fluorescent trace. Significant Ca^2+^ events were then selected with a supervised algorithm as follows. Firstly, a new standard deviation was calculated on the baseline trace, and all local maxima with absolute values, exceeding twice this new standard deviation, were identified. Secondly, among these events, we considered significant only those associated with local Ca^2+^ dynamics with amplitude larger than threefold the new standard deviation. All Ca^2+^ traces were then visually inspected to exclude the ROIs dominated by noise. A Ca^2+^ response was considered due to the electrical stimulation when the Ca^2+^ peak falls in a time-period of 5 s after the stimulus. For each series (LFS1, HFS, LFS2) neurons were classified based on their activity profile: hyperactive (>4 Ca²⁺ events), active (4 Ca²⁺ events), hypoactive (<4 Ca²⁺ events), or silent (no transients during that series, but active in at least one of the others). Neurons with no Ca²⁺ events across all recording series were excluded from the analysis. We then calculated the percentage of responsive neuronal bodies with respect to each single electrical stimulation.

### Statistical Analysis

All statistical analyses were performed using GraphPad Prism 10 or OriginPro 2023b (OriginLab Corporation, Northampton, MA, USA). The statistical test used for each dataset is specified in the corresponding figure legend. Normality was assessed with the Shapiro–Wilk test. For normally distributed data, one-way ANOVA followed by Dunnett’s multiple comparisons test was used to compare each experimental group. For two-group datasets (e.g., WT vs. AD), unpaired two-tailed t-tests were applied when data were normally distributed. For non-normally distributed data, the Kruskal–Wallis test followed by Dunn’s multiple comparisons test was used. A P-value < 0.05 was considered statistically significant. For 2P Ca^2+^ imaging data χ² test followed by pairwise comparisons with Bonferroni correction was used for pie-charts, while Kolmogorov–Smirnov test followed by Bonferroni correction with α = 0.0167 was used for bar histograms.

## Acknowledgements

The research of the authors was funded by grants from the Italian Ministry of University and Scientific Research (PRIN2022943TH9 to P.P., F.D.V., and A.G., and PRIN20225R4Y5 to P.P., financed by the European Union, NextGenerationEU); Cure Alzheimer’s Fund (USA) to P.P., F.D.V., and A.G.; the Alzheimer’s Association (USA; E2A-23-1148250) to P.P.; the Italian Ministry of Research, NRRP–National Recovery and Resilience Plan grant, National Centre of Research “Development of gene therapy and drugs with RNA technology,” CN3 “Neurodegenerative Diseases” to P.P. (NextGenerationEU); and CODBAN_000428, CUP J73C24000090007, Spoke 3 – M4C2 – Inv.1.4 Prog. CN00000041 to F.D.V and A.G.

A.L. research contract was funded by the Italian Ministry of Research, NRRP–National Recovery and Resilience Plan grant, National Centre of Research “Development of gene therapy and drugs with RNA technology,” CN3 “Neurodegenerative Diseases” (NextGenerationEU).

N.K. PhD fellowship was funded by the European Union’s Horizon 2020 research and innovation program under the Marie Sklodowaka-Curie grant agreement no 101034319 and from the European Union – NextGenerationEU.

Imaging instrumentation was made available thanks to: Euro-BioImaging FOE (MUR); National Recovery and Resilience Plan (NRRP), Mission 4 Component 2 Investment 3.1 - Call for tender No. 3264/2021 of Italian MUR, funded by the European Union – NextGenerationEU - Project code IR0000023, Concession Decree No. 101/2022 adopted by the Italian MUR, CUP B53C22001810006, “SEELIFE - Strengthening the Italian Infrastructure of Euro-Bioimaging”.

We are grateful to L. Ozmen and F. Hoffmann-La Roche Ltd (Basel, Switzerland) for kindly donating the AD mice used in this study (MTA 16-02-07-UniPD). We also thank Memmia Srl (Padua, Italy) for generating the AD-P2X7RKO mouse line, and L. Zentilin (Molecular Medicine Laboratory, International Centre for Genetic Engineering and Biotechnology, ICGEB, Trieste, Italy) for producing the AAV used for the delivery of the pmeLUC probe. We also thank Vito Barbieri, technician at the Department of Surgery, Oncology and Gastroenterology, University of Padua, for the organization and invaluable support with the *in vivo* eATP experiments.

Francesco Di Virgilio passed away on September 22, 2024, and made substantial intellectual contributions to this study, including conceptualization, design, supervision, interpretation of results, and funding acquisition. His scientific vision and mentorship profoundly shaped this project. We dedicate this work to his memory. His pioneering research on extracellular ATP and P2X7 receptor in innate and adaptive immunity transformed the field and inspired the entire purinergic scientific community. Francesco was an example of a deeply dedicated and honest scientist, a mentor and a dear friend of many of us.

## Author Contributions

Conceptualization: P.P., F.D.V., E.G., A.L.

Methodology: N.A., A.L., N.R., E.G., D.S., F.D.V.

Investigation: N.A. (staining, imaging, and analysis of microglia morphology, phagocytosis, PNNs and complement proteins), M.B. (2P Ca²⁺ imaging, RT-qPCR, western blot, eATP measurements), N.R. (eATP measurements), A.L. (2P Ca²⁺ imaging, eATP measurements), S.F. and M.T. (RT-qPCR, ELISA), N.K. (staining and imaging of complement proteins and PNNs).

Resources: N.R. (colony management, AD-P2X7RKO line establishment).

Formal analysis: N.A., M.B., N.R., A.L., S.F.

Visualization: A.L., N.K., N.A.

Writing – original draft: E.G., A.L., P.P.

Writing – review & editing: E.G., A.L., P.P., E.B., A.G., D.S.

Supervision: E.G., A.L., P.P., F.D.V., N.R., E.B.

Project administration: E.G., A.L., P.P., F.D.V.

Funding acquisition: P.P., F.D.V., A.G.

## Additional information

The AstroResp code used in this work is publicly available at https://github.com/ladymariot/AstroResp.

## Supplementary information

**Extended Data Figure 1.**
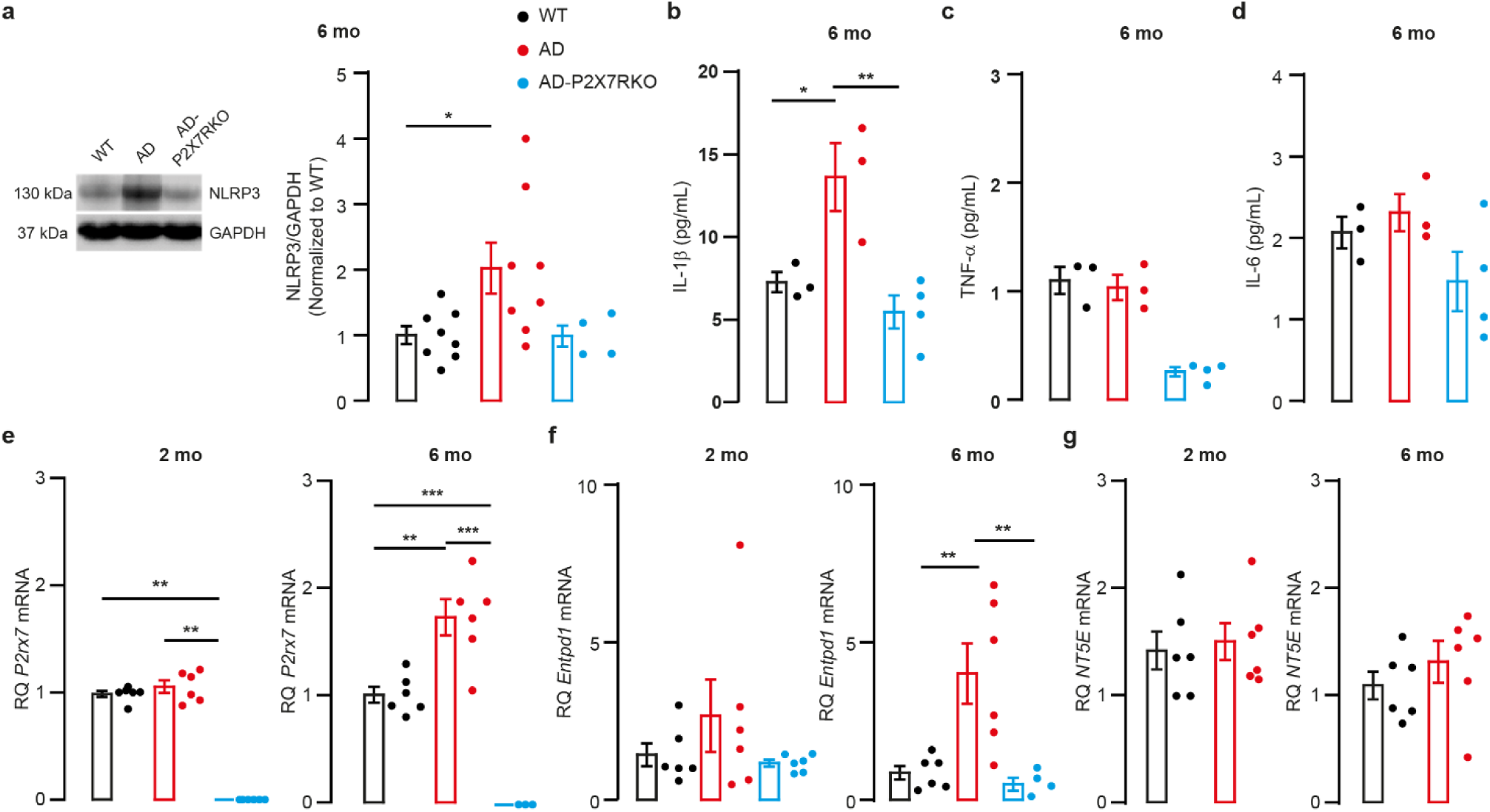
Changes in inflammatory molecules and ectonucleotidases in the mouse cortex during AD progression. **a**, Representative immunoblot of NLRP3 protein, and relative quantification, in cortical lysates from 6-month-old (6 mo) WT (n = 8), AD (n = 8), and AD-P2X7RKO (n = 4) mice. Data are presented as mean ± s.e.m., normalized first to GAPDH and then to WT. **b–d,** Levels of IL-1β (b), TNF-α (c), and IL-6 (d) cytokines in SSCx from 6-month-old WT (n = 3), AD (n = 3), and AD-P2X7RKO (n = 4) mice, measured by ELISA. **e–g**, Quantitative PCR analysis of *P2rx7/*P2X7R (e), *Entpd1*/CD39 (f), and *Nt5e*/CD73 (g) expression in brain cortex from WT, AD, and AD-P2X7RKO mice at 2 mo (WT, n = 6; AD, n = 6; AD-P2X7RKO, n = 6) and 6 mo (WT, n = 6; AD, n = 6; AD-P2X7RKO, n = 3 for *P2rx7,* n = 4 for *Entpd1*). Data are presented as mean ± s.e.m.; statistical analysis was performed using Kruskal–Wallis test followed by Dunn’s post hoc test. For *Nt5e* (g), comparisons between WT and AD were performed using unpaired two-tailed t-tests. \**P* < 0.05; \*\**P* < 0.01; \*\*\**P* < 0.001.

**Extended Data Figure 2.**
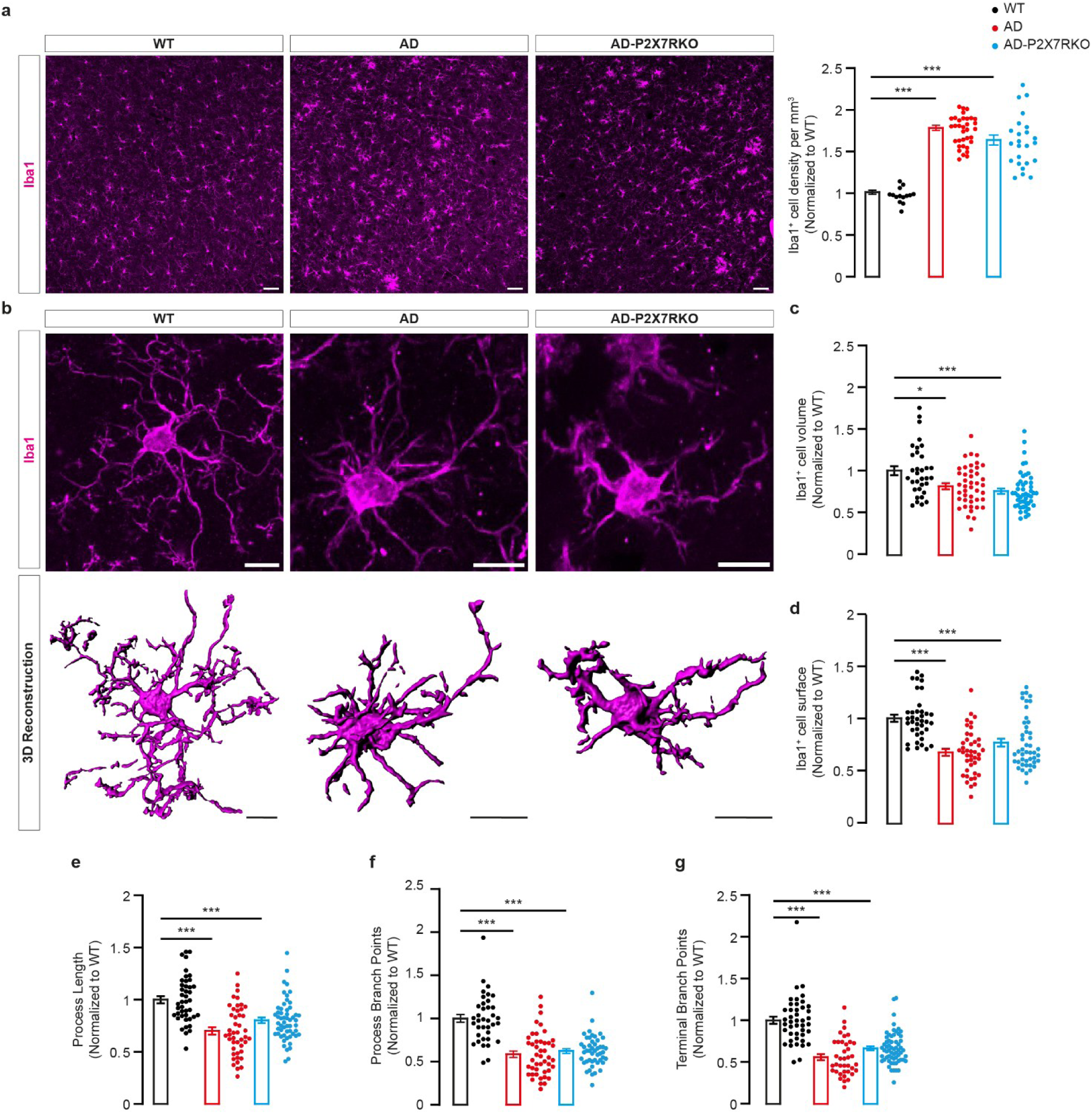
Microglia morphological changes in SSCx L4 of 6-month-old AD mice and their dependency on P2X7R. **a**, Immunofluorescence detection of Iba1⁺ microglia of WT (left), AD (middle), and AD-P2X7RKO (right) mice. Left, representative confocal images acquired at low magnification for microglia density assessment (scale bar, 50 µm); right, quantification of Iba1^+^ cell density in WT (n = 14 FoVs), AD (n = 34 FoVs), and AD-P2X7RKO (n = 24 FoVs) mice. N ≥ 3 animals. **b**, Immunofluorescence detection of Iba1⁺ microglia, as in panel a, imaged at higher magnification for morphological analysis. Top, representative confocal images (scale bar, 10 µm); bottom, 3D reconstructions (scale bar, 10 µm). **c–g**, Quantification of Iba1⁺ microglia volume (c; WT, n = 33 cells; AD, n = 43; AD-P2X7RKO, n = 48), surface area (d; WT, n = 38; AD, n = 42; AD-P2X7RKO, n = 45), process length (e; WT, n = 43; AD, n = 54; AD-P2X7RKO, n = 55), branch points (f; WT, n = 37; AD, n = 45; AD-P2X7RKO, n = 47), and terminal branch points (g; WT, n = 43; AD, n = 38; AD-P2X7RKO, n = 62). N ≥ 3 mice/genotype. Data are presented as mean ± s.e.m. normalized to WT; statistical analysis was performed using Kruskal–Wallis test followed by Dunn’s post hoc test. For panel a and e, one-way ANOVA followed by Dunnett’s multiple comparisons test was applied. \**P* < 0.05; \*\*\**P* < 0.001.

**Extended Data Figure 3.**
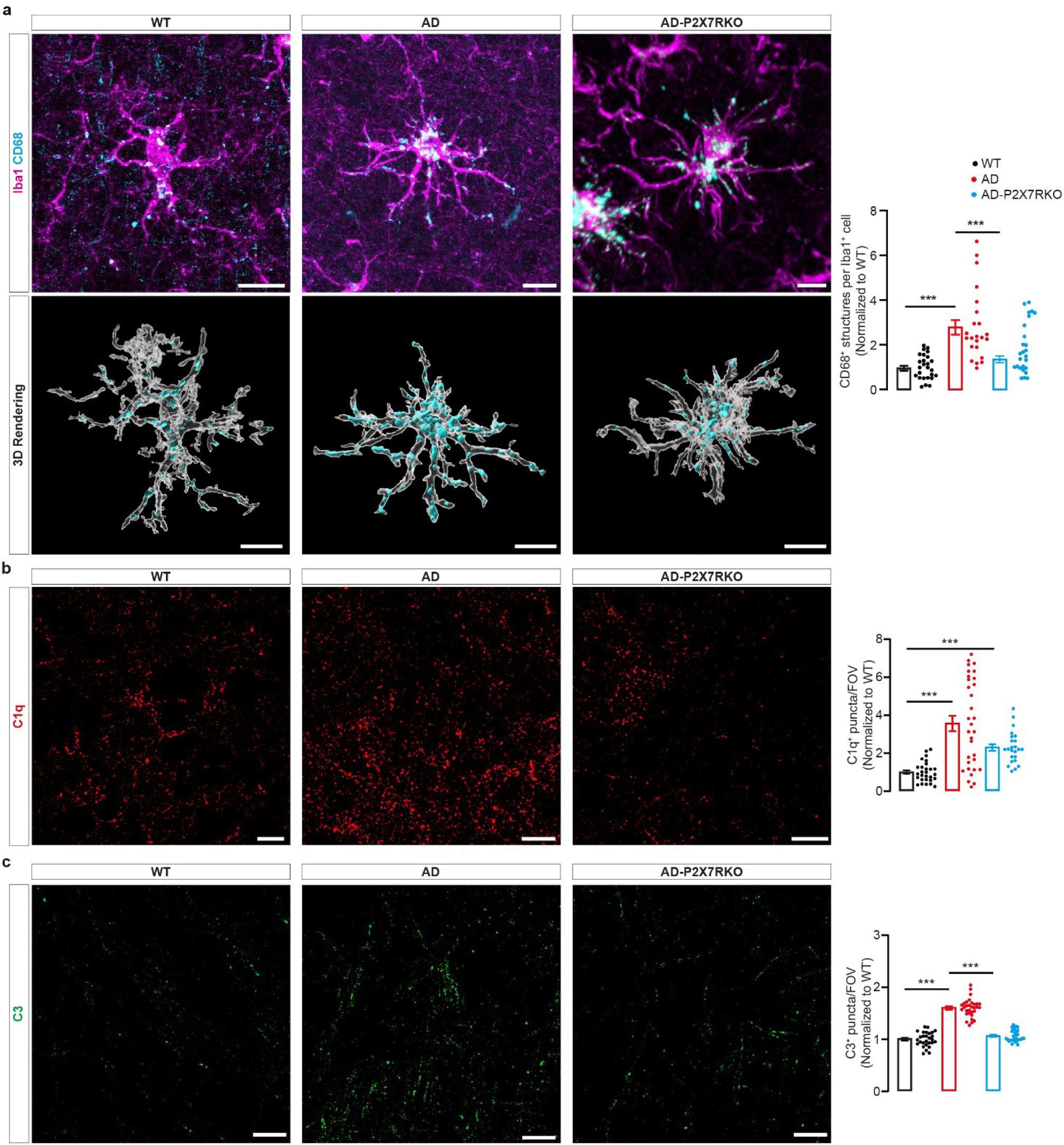
P2X7R drives microglia phagocytic activity and complement activation in SSCx L4 of 6-month-old AD mice. **a** Immunofluorescence detection of CD68⁺ structures within Iba1⁺ microglia of WT (left), AD (middle), and AD-P2X7RKO (right) mice. Top, representative confocal images of microglia immunolabeled for CD68 (cyan) and Iba1 (magenta) (scale bar, 10 µm); bottom, 3D reconstructions (scale bar, 10 µm); right, quantification of CD68⁺ structures per Iba1⁺ cell in WT (n = 58), AD (n = 55), and AD-P2X7RKO (n = 45) mice. N ≥ 3 mice/genotype. **b,** Immunofluorescence detection of C1q⁺ puncta in WT (left), AD (middle), and AD-P2X7RKO (right) mice. Left, representative confocal images (scale bar, 10 µm); right, quantification of C1q^+^ puncta per FoV in 6-month-old WT (n = 28 FoVs), AD (n = 32 FoVs), and AD-P2X7RKO (n = 24 FoVs). N ≥ 3 mice/genotype. **c,** Immunofluorescence detection of C3^+^ puncta in WT (left), AD (middle), and AD-P2X7RKO (right) mice. Left, representative confocal images (scale bar, 10 µm); right, quantification of C3^+^ puncta per FoV in WT (n = 28 FoVs), AD (n = 31 FoVs), and AD-P2X7RKO (n = 31 FoVs) animals. N ≥ 3 mice/genotype. Data are presented as mean ± s.e.m. normalized to WT; statistical analysis was performed using Kruskal–Wallis test followed by Dunn’s post hoc test. \*\*\**P* < 0.001.

**Extended Data Figure 4.**
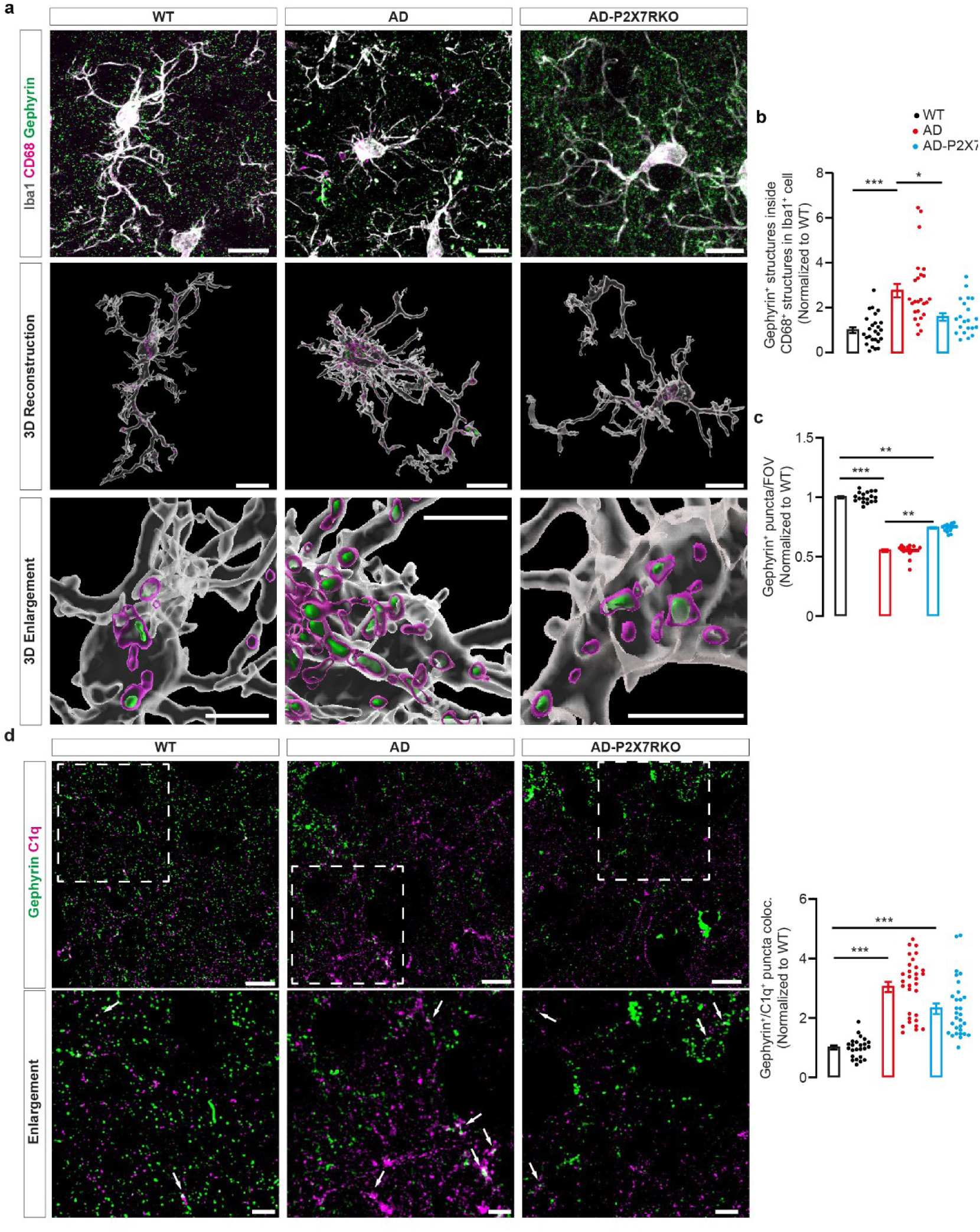
P2X7R KO mitigates complement-mediated engulfment of inhibitory synapses by microglia in SSCx L4 of 6-month-old AD mice. **a,** Immunofluorescence detection of CD68⁺ structures colocalized with gephyrin^+^ synapses within Iba1⁺ microglia in WT (left), AD (middle), and AD-P2X7RKO (right) mice. Top, representative confocal images of Iba1⁺ (grey) microglia also immunolabeled for CD68 (magenta) and gephyrin (green) (scale bar, 10 µm); middle, 3D reconstructions (scale bar, 10 µm); bottom, 3D reconstruction enlargement (scale bar, 5 µm). **b**, Quantification of CD68⁺ structures colocalized with gephyrin^+^ puncta per Iba1⁺ cell in WT (n = 27 FoVs), AD (n = 25 FoVs), and AD-P2X7RKO (n = 21 FoVs) mice. N ≥ 3 mice/genotype. **c**, Quantification of gephyrin^+^ puncta per FOV in WT (n = 18 FoVs), AD (n = 19 FoVs), and AD-P2X7RKO (n = 17 FoVs) mice. N ≥ 3 mice/genotype. **d**, Immunofluorescence detection of C1q⁺ puncta colocalized with gephyrin⁺ synapses in WT (left), AD (middle), and AD-P2X7RKO (right) mice. Top left, representative confocal images of C1q⁺ (magenta) and gephyrin⁺ (green) puncta (scale bar, 5 µm), with at the bottom the enlargement on the dashed box, white arrows indicate representative colocalizations (scale bar, 2 µm); right, quantification of C1q⁺ puncta colocalized with gephyrin⁺ ones in WT (n = 24 FoVs), AD (n = 31 FoVs), and AD-P2X7RKO (n = 30 FoVs) mice. N ≥ 3 mice/genotype. Data are presented as mean ± s.e.m. normalized to WT; statistical analysis was performed using Kruskal–Wallis test followed by Dunn’s post hoc test. \**P* < 0.05; ** *P* < 0.01; *** *P* < 0.001.

**Extended Data Figure 5.**
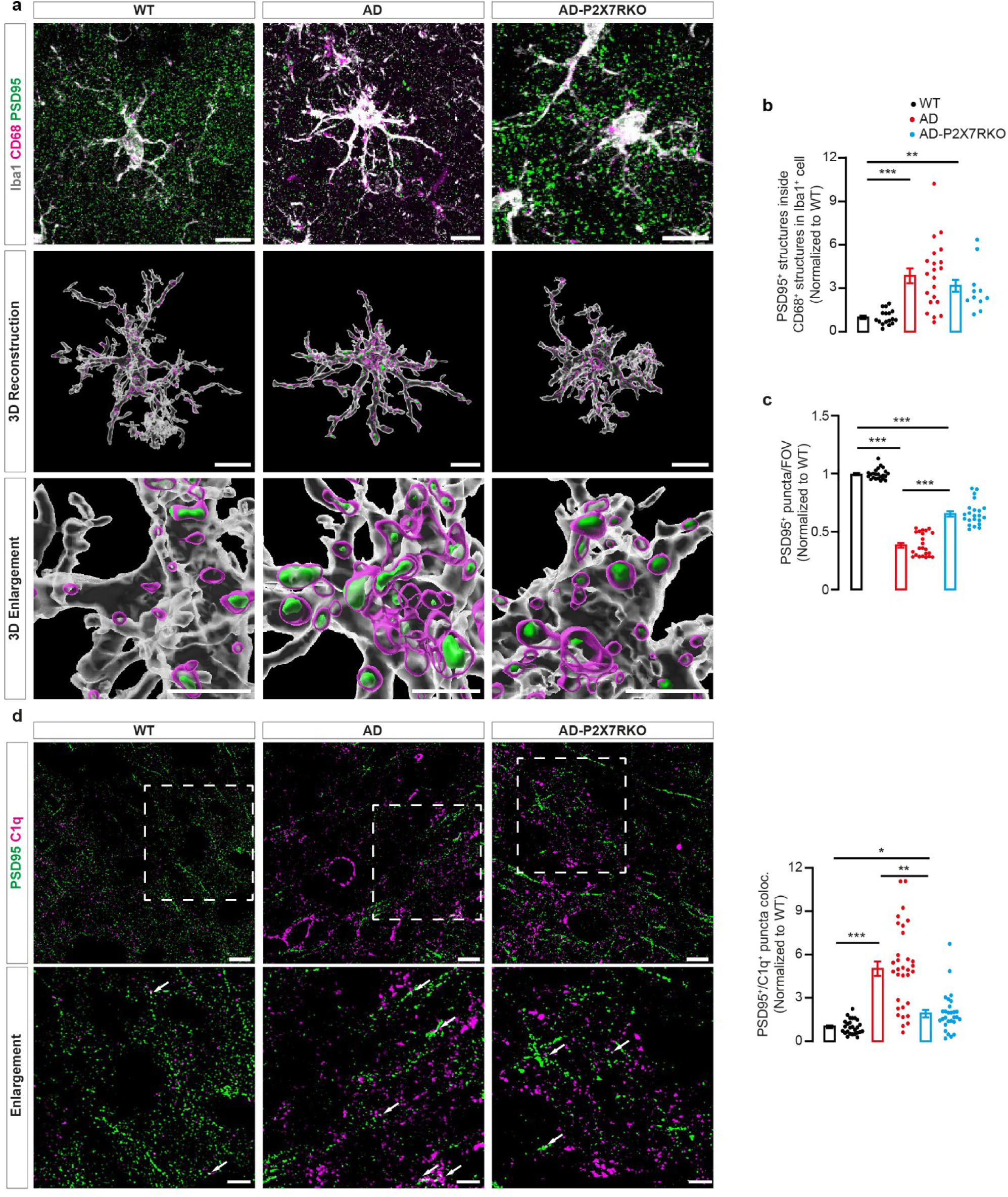
P2X7R KO mitigates complement-mediated engulfment of excitatory synapses by microglia in SSCx L4 of 6-month-old AD mice. **a,** Immunofluorescence detection of CD68⁺ structures colocalized with PSD95^+^ synapses within Iba1⁺ microglia in WT (left), AD (middle), and AD-P2X7RKO (right) mice. Top, representative confocal images of Iba1⁺ (white) microglia also immunolabeled for CD68 (magenta) and PSD95 (green) (scale bar, 10 µm); middle, 3D reconstructions (scale bar, 10 µm); bottom, 3D reconstruction enlargement (scale bar, 5 µm). **b**, Quantification of CD68⁺ structures colocalized with PSD95^+^ puncta per Iba1⁺ cell in WT (n = 17 FoVs), AD (n = 21 FoVs), and AD-P2X7RKO (n = 14 FoVs) mice. N ≥ 3 mice/genotype. **c**, Quantification of PSD95^+^ puncta per FoV in WT (n = 23 FoVs), AD (n = 23 FoVs), and AD-P2X7RKO (n = 22 FoVs) mice. N ≥ 3 mice/genotype. **d**, Immunofluorescence detection of C1q⁺ puncta colocalized with PSD95⁺ synapses in WT (left), AD (middle), and AD-P2X7RKO (right) mice. Top left, representative confocal images of C1q⁺ (magenta) and PSD95⁺ (green) puncta (scale bar, 5 µm), with at the bottom the enlargement on the dashed box, white arrows indicate representative colocalizations (scale bar, 2 µm); right, quantification of C1q⁺ puncta colocalized with PSD95⁺ ones in WT (n = 26 FoVs), AD (n = 32 FoVs), and AD-P2X7RKO (n = 28 FoVs) mice. N ≥ 3 mice/genotype. Data are presented as mean ± s.e.m. normalized to WT; statistical analysis was performed using Kruskal–Wallis test followed by Dunn’s post hoc test. In panel f, one-way ANOVA followed by Dunnett’s multiple comparisons test was applied. \**P* < 0.05; \*\**P* < 0.01; \*\*\**P* < 0.001.

**Extended Data Figure 6.**
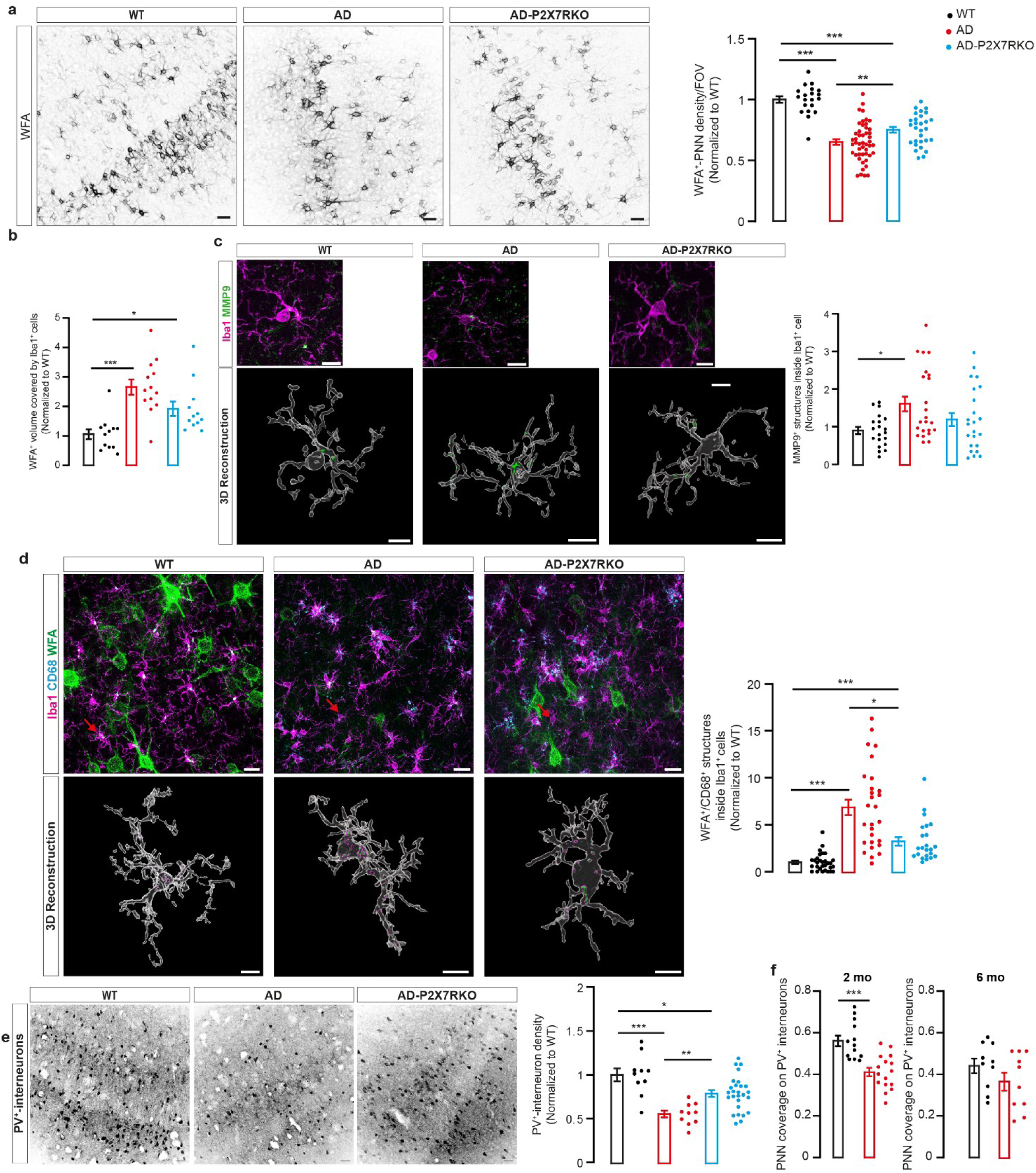
P2X7R KO mitigates microglia-mediated PNN remodeling and interneuron vulnerability in SSCx L4 of 6-month-old AD mice. **a**, Left, WFA⁺-PNN structures in WT (left), AD (middle), and AD-P2X7RKO (right) mice. Representative confocal images (scale bar, 50 µm). Right, quantification of WFA⁺ structures per FoV in WT (n = 20 FoVs), AD (n = 48 FoVs), and AD-P2X7RKO (n = 29 FoVs) mice. N ≥ 3 mice/genotype. **b**, Quantification of WFA⁺-PNN structures colocalized with Iba1⁺ cells in WT (n = 12 FoVs), AD (n = 13 FoVs), and AD-P2X7RKO (n = 12 FoVs) mice. N ≥ 3 mice/genotype. **c**, Left, immunofluorescence detection of MMP9⁺ structures within Iba1⁺ microglia in WT (left), AD (middle), and AD-P2X7RKO (right) mice. Top, representative confocal images of Iba1⁺ (magenta) microglia also immunolabeled for MMP9 (green, scale bar, 10 µm); bottom, 3D reconstructions (scale bar, 10 µm). Right, quantification of MMP9⁺ structures within Iba1⁺ cells in WT (n = 20 FoVs), AD (n = 23 FoVs), and AD-P2X7RKO (n = 23 FoVs) mice. N ≥ 3 mice/genotype. **d**, Left, Immunofluorescence detection of CD68⁺ structures colocalized with WFA⁺-PNN structures within Iba1⁺ microglia in WT (left), AD (middle), and AD-P2X7RKO (right) mice. Top, representative confocal images of Iba1⁺ (magenta) microglia also labeled for CD68 (cyan) and WFA⁺ structures (green, scale bar, 20 µm). Red arrows indicate the reconstructed microglial cell; bottom, 3D reconstructions (scale bar, 10 µm). Right, quantification of WFA⁺-PNN structures colocalized with CD68⁺ structures within Iba1⁺ cells in WT (n = 30 FoVs), AD (n = 28 FoVs), and AD-P2X7RKO (n = 23 FoVs) mice. N ≥ 3 mice/genotype. **e**, Left, representative confocal images of PV⁺ interneurons in WT (left), AD (middle), and AD-P2X7RKO (right) mice (scale bar, 50 µm). Right, quantification of PV⁺ interneuron density per FoV in WT (n = 10 FoVs), AD (n = 10 FoVs), and AD-P2X7RKO (n = 27 FoVs) mice. N ≥ 3 mice/genotype. **f**, Quantification of PV⁺ neurons covered by WFA⁺-PNN structures per FOV in 2 or 6-month-old WT (n = 13 and 10 FoVs for 2 and 6-month-old WT mice respectively) and AD (n = 16 and 10 FoVs for 2 and 6-month-old WT mice respectively). N ≥ 3 mice/genotype. Data are presented as mean ± s.e.m. normalized to WT; statistical analysis was performed using Kruskal–Wallis test followed by Dunn’s post hoc test. For Panel a, d and e one-way ANOVA followed by Dunnett’s multiple comparisons test was applied. \**P* < 0.05; \*\**P* < 0.01; \*\*\**P* < 0.001.

## References

1. Henstridge, C. M., Hyman, B. T. & Spires-Jones, T. L. Beyond the neuron–cellular interactions early in Alzheimer disease pathogenesis. Nat Rev Neurosci 20, 94–108 (2019).

2. Sierra, A., Miron, V. E., Paolicelli, R. C. & Ransohoff, R. M. Microglia in Health and Diseases: Integrative Hubs of the Central Nervous System (CNS). Cold Spring Harb Perspect Biol 16, a041366 (2024).

3. Tewari, B. P., Chaunsali, L., Prim, C. E. & Sontheimer, H. A glial perspective on the extracellular matrix and perineuronal net remodeling in the central nervous system. Front Cell Neurosci 16, (2022).

4. Tsering, W. et al. Preferential clustering of microglia and astrocytes around neuritic plaques during progression of Alzheimer’s disease neuropathological changes. J Neurochem 169, (2025).

5. Leng, F. & Edison, P. Neuroinflammation and microglial activation in Alzheimer disease: where do we go from here? Nat Rev Neurol 17, 157–172 (2021).

6. Awogbindin, I., Wanklin, M., Verkhratsky, A. & Tremblay, M.-È. Microglia in Neurodegenerative Diseases. in 497–512 (2024). doi:10.1007/978-3-031-55529-9_27.

7. Di Virgilio, F., Vultaggio-Poma, V., Falzoni, S. & Giuliani, A. L. Extracellular ATP: A powerful inflammatory mediator in the central nervous system. Neuropharmacology 224, 109333 (2023).

8. Giuliani, A. L., Sarti, A. C. & Di Virgilio, F. Ectonucleotidases in Acute and Chronic Inflammation. Front Pharmacol 11, (2021).

9. Badimon, A. et al. Negative feedback control of neuronal activity by microglia. Nature 586, 417–423 (2020).

10. Liu, Y. et al. Research progress on adenosine in central nervous system diseases. CNS Neurosci Ther 25, 899–910 (2019).

11. Di Virgilio, F., Dal Ben, D., Sarti, A. C., Giuliani, A. L. & Falzoni, S. The P2X7 Receptor in Infection and Inflammation. Immunity 47, 15–31 (2017).

12. Martin, E. et al. New role of P2X7 receptor in an Alzheimer’s disease mouse model. Mol Psychiatry 24, 108–125 (2019).

13. McLarnon, J. G., Ryu, J. K., Walker, D. G. & Choi, H. B. Upregulated expression of purinergic P2X7 receptor in Alzheimer disease and amyloid-β peptide-treated microglia and in peptide-injected rat hippocampus. J Neuropathol Exp Neurol. 2006 Nov;65(11):1090–7.

14. Huang, Q. et al. P2X7 Receptor: an Emerging Target in Alzheimer’s Disease. Mol Neurobiol 61, 2866–2880 (2024).

15. Ozmen, L., Albientz, A., Czech, C. & Jacobsen, H. Expression of Transgenic APP mRNA Is the Key Determinant for Beta-Amyloid Deposition in PS2APP Transgenic Mice. Neurodegener Dis 6, 29–36 (2009).

16. Pellegatti, P., Falzoni, S., Pinton, P., Rizzuto, R. & Di Virgilio, F. A Novel Recombinant Plasma Membrane-targeted Luciferase Reveals a New Pathway for ATP Secretion. Mol Biol Cell 16, 3659–3665 (2005).

17. Lia, A. et al. Rescue of astrocyte activity by the calcium sensor STIM1 restores long-term synaptic plasticity in female mice modelling Alzheimer’s disease. Nat Commun 14, 1590 (2023).

18. Sim, J. A., Young, M. T., Sung, H.-Y., North, R. A. & Surprenant, A. Reanalysis of P2X _7_ Receptor Expression in Rodent Brain. The Journal of Neuroscience 24, 6307–6314 (2004).

19. Savage JC, Carrier M, Tremblay MÈ. Morphology of Microglia Across Contexts of Health and Disease. Methods Mol Biol. 2019;2034:13–26.

20. Paolicelli, R. C. et al. Microglia states and nomenclature: A field at its crossroads. Neuron 110, 3458–3483 (2022).

21. Miao, J. et al. Microglia in Alzheimer’s disease: pathogenesis, mechanisms, and therapeutic potentials. Front Aging Neurosci 15, (2023).

22. Scott-Hewitt, N., Huang, Y. & Stevens, B. Convergent mechanisms of microglia-mediated synaptic dysfunction contribute to diverse neuropathological conditions. Ann N Y Acad Sci 1525, 5–27 (2023).

23. Taddei, R. N. & Duff, K. E. Synapse vulnerability and resilience across the clinical spectrum of dementias. Nat Rev Neurol 21, 353–369 (2025).

24. Ayata, P. et al. Epigenetic regulation of brain region-specific microglia clearance activity. Nat Neurosci 21, 1049–1060 (2018).

25. Stevens, B. et al. The Classical Complement Cascade Mediates CNS Synapse Elimination. Cell 131, 1164–1178 (2007).

26. Totaro, V., Pizzorusso, T. & Tognini, P. Orchestrating the Matrix: The Role of Glial Cells and Systemic Signals in Perineuronal Net Dynamics. Neurochem Res 50, 253 (2025).

27. Auer, S. et al. The Role of Perineuronal Nets in Physiology and Disease: Insights from Recent Studies. Cells 14, 321 (2025).

28. Crapser, J. D., Arreola, M. A., Tsourmas, K. I. & Green, K. N. Microglia as hackers of the matrix: sculpting synapses and the extracellular space. Cell Mol Immunol 18, 2472–2488 (2021).

29. Lorenzl, S. Increased plasma levels of matrix metalloproteinase-9 in patients with Alzheimer’s disease. Neurochem Int 43, 191–196 (2003).

30. Radosinska, D. & Radosinska, J. The Link Between Matrix Metalloproteinases and Alzheimer’s Disease Pathophysiology. Mol Neurobiol 62, 885–899 (2025).

31. Targa Dias Anastacio, H., Matosin, N. & Ooi, L. Neuronal hyperexcitability in Alzheimer’s disease: what are the drivers behind this aberrant phenotype? Transl Psychiatry 12, 257 (2022).

32. Citri, A. & Malenka, R. C. Synaptic Plasticity: Multiple Forms, Functions, and Mechanisms. Neuropsychopharmacology 33, 18–41 (2008).

33. Heneka, M. T. et al. Neuroinflammation in Alzheimer disease. Nat Rev Immunol 25, 321–352 (2025).

34. Faroqi, A. H. et al. *In Vivo* Detection of Extracellular Adenosine Triphosphate in a Mouse Model of Traumatic Brain Injury. J Neurotrauma 38, 655–664 (2021).

35. Calovi, S., Mut-Arbona, P. & Sperlágh, B. Microglia and the Purinergic Signaling System. Neuroscience 405, 137–147 (2019).

36. Abdelhamed, H. G. et al. Brain interleukins and Alzheimer’s disease. Metab Brain Dis 40, 116 (2025).

37. Bedetta, M., Pizzo, P. & Lia, A. The Multifaceted Role of P2X7R in Microglia and Astrocytes. Neurochem Res 50, 239 (2025).

38. Mathys, H. et al. Single-cell transcriptomic analysis of Alzheimer’s disease. Nature 570, 332–337 (2019).

39. Mathys, H. et al. Temporal Tracking of Microglia Activation in Neurodegeneration at Single-Cell Resolution. Cell Rep 21, 366–380 (2017).

40. Mei, S.-Y., Zhang, N., Wang, M., Lv, P. & Liu, Q. Microglial purinergic signaling in Alzheimer’s disease. Purinergic Signal 21, 815–827 (2025).

41. Robel, S. & Sontheimer, H. Glia as drivers of abnormal neuronal activity. Nat Neurosci 19, 28–33 (2016).

42. Yang, H., Andersson, U. & Brines, M. Neurons Are a Primary Driver of Inflammation via Release of HMGB1. Cells 10, 2791 (2021).

43. Pawelec, P., Ziemka-Nalecz, M., Sypecka, J. & Zalewska, T. The Impact of the CX3CL1/CX3CR1 Axis in Neurological Disorders. Cells 9, 2277 (2020).

44. Zhang, X. et al. Epigenetic regulation of innate immune memory in microglia. J Neuroinflammation 19, 111 (2022).

45. Hodebourg, R., Scofield, M. D., Kalivas, P. W. & Kuhn, B. N. Nonneuronal contributions to synaptic function. Neuron 113, 2399–2415 (2025).

46. Wen, L., Bi, D. & Shen, Y. Complement-mediated synapse loss in Alzheimer’s disease: mechanisms and involvement of risk factors. Trends Neurosci 47, 135–149 (2024).

47. Lui, H. et al. Progranulin Deficiency Promotes Circuit-Specific Synaptic Pruning by Microglia via Complement Activation. Cell 165, 921–935 (2016).

48. Dejanovic, B. et al. Complement C1q-dependent excitatory and inhibitory synapse elimination by astrocytes and microglia in Alzheimer’s disease mouse models. Nat Aging 2, 837–850 (2022).

49. Hong, S. et al. Complement and microglia mediate early synapse loss in Alzheimer mouse models. Science (1979) 352, 712–716 (2016).

50. Werneburg, S. et al. Targeted Complement Inhibition at Synapses Prevents Microglial Synaptic Engulfment and Synapse Loss in Demyelinating Disease. Immunity 52, 167–182.e7 (2020).

51. Zhang, W., Chen, Y. & Pei, H. C1q and central nervous system disorders. Front Immunol 14, (2023).

52. Chen, Y., Chu, J. M.-T., Wong, G. T.-C. & Chang, R. C.-C. Complement C3 From Astrocytes Plays Significant Roles in Sustained Activation of Microglia and Cognitive Dysfunctions Triggered by Systemic Inflammation After Laparotomy in Adult Male Mice. Journal of Neuroimmune Pharmacology 19, 8 (2024).

53. Chen, Z.-P. et al. GABA-dependent microglial elimination of inhibitory synapses underlies neuronal hyperexcitability in epilepsy. Nat Neurosci 28, 1404–1417 (2025).

54. Crapser, J. D. et al. Microglia facilitate loss of perineuronal nets in the Alzheimer’s disease brain. EBioMedicine 58, 102919 (2020).

55. Hijazi, S. et al. Early restoration of parvalbumin interneuron activity prevents memory loss and network hyperexcitability in a mouse model of Alzheimer’s disease. Mol Psychiatry 25, 3380–3398 (2020).

56. Allami, P., Yazdanpanah, N. & Rezaei, N. The role of neuroinflammation in PV interneuron impairments in brain networks; implications for cognitive disorders. Rev Neurosci 36, 497–517 (2025).

57. Leparulo, A. et al. Dampened Slow Oscillation Connectivity Anticipates Amyloid Deposition in the PS2APP Mouse Model of Alzheimer’s Disease. Cells 9, 54 (2019).

58. Fontana, R. et al. Early hippocampal hyperexcitability in PS2APP mice: role of mutant PS2 and APP. Neurobiol Aging 50, 64–76 (2017).

59. Kipanyula, M. J. et al. Ca ^2+^ dysregulation in neurons from transgenic mice expressing mutant presenilin 2. Aging Cell 11, 885–893 (2012).

60. Engel, T. et al. Seizure suppression and neuroprotection by targeting the purinergic P2X7 receptor during status epilepticus in mice. The FASEB Journal 26, 1616–1628 (2012).

61. Schmidt, S., Isaak, A. & Junker, A. Spotlight on P2X7 Receptor PET Imaging: A Bright Target or a Failing Star? Int J Mol Sci 24, 1374 (2023).

62. Qiu, L. et al. Synthesis and in vitro evaluation of novel compounds and discovery of a promising iodine-125 radioligand for purinergic P2X7 receptor (P2X7R). Bioorg Med Chem 118, 118054 (2025).

63. Apolloni, S. et al. Novel P2X7 Antagonist Ameliorates the Early Phase of ALS Disease and Decreases Inflammation and Autophagy in SOD1-G93A Mouse Model. Int J Mol Sci 22, 10649 (2021).

64. Eli Lilly and Asahi Kasei Pharma. Press release: License agreement for chronic pain drug candidate. (2024). Available at: https://investor.lilly.com/news-releases/news-release-details/lilly-and-asahi-kasei-pharma-announce-license-agreement-chronic.

## References

1. Ozmen, L., Albientz, A., Czech, C. & Jacobsen, H. Expression of Transgenic APP mRNA Is the Key Determinant for Beta-Amyloid Deposition in PS2APP Transgenic Mice. Neurodegener Dis 6, 29–36 (2009).

2. Sim, J. A., Young, M. T., Sung, H.-Y., North, R. A. & Surprenant, A. Reanalysis of P2X 7 Receptor Expression in Rodent Brain. The Journal of Neuroscience 24, 6307–6314 (2004).

3. Dejanovic, B. et al. Complement C1q-dependent excitatory and inhibitory synapse elimination by astrocytes and microglia in Alzheimer’s disease mouse models. Nat Aging 2, 837–850 (2022).

4. Lia, A. et al. Rescue of astrocyte activity by the calcium sensor STIM1 restores long-term synaptic plasticity in female mice modelling Alzheimer’s disease. Nat Commun 14, 1590 (2023).

